# Allosteric network in Ube2T drives specificity for RING E3 catalysed ubiquitin signals

**DOI:** 10.1101/429076

**Authors:** Viduth K Chaugule, Connor Arkinson, Rachel Toth, Helen Walden

## Abstract

In eukaryotes, DNA damage repair is implemented by a host of proteins that are coordinated by defined molecular signals. One such signal that transpires during the Fanconi Anemia (FA) - interstrand crosslink (ICL) repair pathway is the site-specific monoubiquitination of FANCD2 and FANCI proteins by a large, multi-protein FA core complex. The mechanics for this exquisitely specific monoubiquitin signal has been elusive. Here we show FANCL, the RING E3 module of the FA core complex, allosterically activates its cognate E2 Ube2T for monoubiquitination by a mechanism distinct from the typical RING-based catalysis. FANCL triggers intricate re-wiring of Ube2T’s intra-residue network thus activating the E2 for precision targeting. This network is intrinsically regulated by conserved gates and loops which can be engineered to yield Ube2T variants that enhance FANCD2 ubiquitination by ~30-fold without compromising on target specificity. Finally, we also uncover allosteric networks in other ubiquitin E2s that can be leveraged by RING E3 ligases to drive specific ubiquitination.

## Introduction

Ubiquitination is an essential, versatile and reversible post-translational modification system that enables eukaryotic cells to reprogram the fate and function of the modified protein and its connected pathway. The modification is accomplished by a sequential enzyme cascade wherein ubiquitin’s C-terminus is first activated by an E1, is transferred onto the catalytic cysteine of an E2 conjugase (E2~Ub) and finally E3 ligases mediate the covalent attachment of ubiquitin onto a target lysine residue (Hochstrasser, 2009; Pickart, 2001). The Really Interesting New Gene (RING) ligase proteins represent the largest E3 family (~600 members) which share a zinc coordinating cross-brace motif termed RING domain (Freemont, 2000). Mechanistically, while non-RING elements of E3 ligases specify the substrate, the RING domains bind an E2 surface distal from the E2 active site and indirectly induce substrate ubiquitination by stabilizing a productive E2~Ub conformation (Metzger et al., 2014). Moreover, any of the seven surface lysine residues on ubiquitin or its N-terminus can be targeted to build polyubiquitin chains, thus enabling diverse signals (Kulathu and Komander, 2012). Typically, RING-E2 interactions are found to be transient thus allowing E3s to switch their E2 partners in order to assemble polyubiquitin signals on the substrate (Brown et al., 2014; Kelly et al., 2014; Rodrigo-Brenni and Morgan, 2007; Windheim et al., 2008). Around 35 ubiquitin E2s are found in mammals, several of which build chains (Stewart et al., 2016). Mechanisms of chain-assembly are well understood and generally involve additional non-covalent interactions between the E2 and the acceptor ubiquitin surface proximal to the linkage site (Eddins et al., 2006; Liu et al., 2014; Middleton and Day, 2015; Petroski and Deshaies, 2005; Rodrigo-Brenni et al., 2010; Wickliffe et al., 2011). In contrast however, far less is known about how RING E3-E2 enzyme pairs attach a single ubiquitin directly on the substrate surface in the case of monoubiquitination.

Site-specific monoubiquitin signals feature prominently in fundamental DNA damage response pathways (Al-Hakim et al., 2010; Uckelmann and Sixma, 2017). In eukaryotes, the repair of toxic DNA interstrand cross-links (ICL) is mediated by the Fanconi Anemia (FA) pathway, defects in which give rise to FA, a genome instability disorder typified by bone marrow failure and high predisposition to cancers (Garaycoechea and Patel, 2014; Kottemann and Smogorzewska, 2013). A key event in FA-ICL damage response is the site-specific mono-ubiquitination of two large (~160kDa) structurally homologous proteins, FANCD2 (Garcia-Higuera et al., 2001) and FANCI (Sims et al., 2007; Smogorzewska et al., 2007) (Lys561 and Lys523 respectively in humans), that signals the recruitment of repair factors (Ceccaldi et al., 2016). The specific modification is mediated by the RING bearing protein FANCL, present within a nine-protein FA core-complex E3 ligase (FANCA-FANCG-FAAP20-FANCC-FANCE-FANCF-FANCB-FANCL-FAAP100) and the E2 Ube2T (Machida et al., 2006; Meetei et al., 2003; Walden and Deans, 2014). FANCL’s central region, a bi-lobed UBC (Ubiquitin conjugation fold)-RWD domain, facilitates direct FANCD2/FANCI interaction (Cole et al., 2010; Hodson et al., 2011) while the C-terminal RING domain selectively binds Ube2T over other E2’s (Hodson et al., 2014). Further, genetic mutations in Ube2T, a Class III E2 with an unstructured C-terminal extension, have recently been linked to a FA phenotype (Hira et al., 2015; Rickman et al., 2015; Virts et al., 2015). Accordingly, in *in vitro* assays, the isolated FANCL and Ube2T enzymes catalyse FANCD2 monoubiquitination, although the modification levels are unexpectedly low (Alpi et al., 2008; Hodson et al., 2014). Studies in frog egg extracts show majority of FANCD2 is present in complex with FANCI (Sareen et al., 2012) while the cell based data indicate mono-ubiquitination of either protein requires the presence of the partner (Alpi and Patel, 2009). However, a crystal structure of the mouse FANCI-FANCD2 complex reveals an extended heterodimer interface which buries the respective target lysine (Joo et al., 2011). Notably, the addition of structured or duplex DNA greatly stimulates FANCD2 ubiquitination and requires the presence of FANCI (Longerich et al., 2014; Sato et al., 2012). The DNA binding propensity of the FANCI-FANCD2 complex, absent in FANCL or Ube2T, is proposed to induce local reconfiguration that could improve FANCL’s access to the target sites. Finally, while FANCL alone induces low levels of ubiquitination, however when in a FANCB-FANCL-FAAP100 sub-complex FANCD2 monoubiquitination levels improve by around 5-fold. The added presence of a FANCC-FANCE-FANCF sub-complex progressively enhances the tandem mono-ubiquitination of the FANCD2-FANCI complex (Rajendra et al., 2014; van Twest et al., 2017). Thus, in the current model the presence of DNA, FANCI and additional FA sub-complexes are all required for the modification however, underlying mechanisms for the sub-complex induced enhancement are not well understood. First, FANCL dependent FANCD2 ubiquitination has been observed in non-vertebrate species which lack an intact FA core-complex, suggesting in part that the mechanism for the specific ubiquitination is encoded within FANCL (Sugahara et al., 2012; Zhang et al., 2009). Second, global proteomic profiling have uncovered several other lysine on human FANCD2 (22 sites) and FANCI (44 sites) that are ubiquitinated *in vivo* indicating the surface of both proteins are viable acceptors (Kim et al., 2011; Udeshi et al., 2013). As ubiquitin signalling is influential in almost every cellular process in eukaryotes, a major unanswered question is how specific signals are assembled and regulated. The FA-ICL repair pathway is crucial for cellular homeostasis thus, understanding how FANCL targets precise FANCD2 and FANCI sites for strict monoubiquitination would provide valuable insights into the mechanics of this crucial DNA damage response signal as well as how target and signal specificity is achieved in ubiquitination.

In this study we show that FANCL activates Ube2T for ubiquitination through an allosteric mechanism that is distinct from the typical RING E3 based catalysis. We find that in addition to the selective FANCL RING-Ube2T interface, there are multiple E2-E3 interactions that perturb the resting state of Ube2T to induce activity. Residue network analysis reveals subtle reconfigurations of Ube2T’s internal connections that link the effect of FANCL binding to the E2’s catalytic centre culminating in substrate ubiquitination. We further uncover intrinsic regulation of this network by conserved Ube2T residues, and through rationally designed mutations we can enhance FANCL mediated FANCD2 (~30 fold) and FANCI (~16 fold) mono-ubiquitination without compromising its specificity. Finally, we identify similar allosteric networks in other ubiquitin E2s that are appropriated by RING E3 ligases to drive specific ubiquitination events.

## Results

### The E2 – E3 pair Ube2T – FANCL ubiquitinates its substrates via an atypical mechanism

Previous *in vitro* studies using recombinant chicken (Alpi et al., 2008; Sato et al., 2012), frog (Hodson et al., 2014) and human (Longerich et al., 2014) proteins report the isolated FANCL enzyme with Ube2T directs FANCD2 monoubiquitination at its physiological target site. However, the underlying mechanism for the site-specific and strict mono-modification is unclear. In order to understand how the Ube2T and FANCL enzyme pair catalyse this specific signal we reconstituted a minimal E2 – E3 module with human proteins. Based on our earlier work we designed and purified a FANCL URD-RING fragment (FANCL^UR^, residues 109-375) that is stable, monomeric (Supplementary Fig 1A) and comprises both the substrate (UBC-RWD domain) and the E2 (RING domain) binding regions (Hodson et al., 2011; Hodson et al., 2014). We then tested the activity of the FANCL^UR^ fragment in *in vitro* FANCD2 ubiquitination assays using fluorescently labelled ubiquitin (Ub^IR800^). Previous studies have shown the additional requirements of FANCI and DNA in complex with FANCD2 for efficient monoubiquitination of this substrate. Consistent with this we observe Ube2T – FANCL^UR^ mediated FANCD2 modification when present as a FANCD2-FANCI-dsDNA complex (Fig 1A). Further, an arginine mutant of the physiological FANCD2 target site (FANCD2 K561R) prevents ubiquitination thus confirming the minimal E2 – E3 module is both active and site-specific. We wondered if the FANCL^UR^ fragment could also specifically ubiquitinate FANCI present in the FANCD2-FANCI-dsDNA complex. To test this we titrated increasing amounts of the E2 – E3 module in reactions containing single target site complexes (FANCI^K523R^-FANCD2 or FANCI-FANCD2^K561R^). FANCL^UR^ clearly favours modification of the FANCD2 site (Fig 1B). In contrast, FANCL^UR^ can drive the site-specific ubiquitination of an isolated FANCI-dsDNA complex (Fig 1C), as previously observed (Longerich et al., 2014). Therefore, unless otherwise stated, all FANCD2 assays are in the presence of FANCI and dsDNA, while FANCI assays are in the presence of dsDNA. RING E3 ligases catalyse ubiquitination by activating the thioester linked E2~ubiquitin (Ub) intermediate. In brief, RING binding of E2~Ub induces the thioester linked Ub to fold back over the E2 wherein the Ile44-centered hydrophobic patch of ubiquitin packs against a central E2 helix. Concomitantly, a ‘linchpin’ RING E3 residue (usually Arg/Lys) contacts both E2 and Ub to stabilise the ‘closed’ E2~Ub conformer, thus priming the thioester for lysine attack (Dou et al., 2012; Plechanovova et al., 2012; Pruneda et al., 2012; Saha et al., 2011). Interestingly, FANCL lacks this linchpin harbouring instead a serine residue (S363) at the analogous position (Supplementary Fig 1B). We therefore wondered if in the absence of a linchpin residue the Ub Ile44 patch requirement is maintained for FANCL’s E3 activity. To test this we compared FANCI and FANCD2 ubiquitination activity of the Ube2T – FANCL^UR^ pair with the well characterised E2 – E3 pair Ube2D3 – RNF4^RING-RING^ (Branigan et al., 2015; Plechanovova et al., 2011; Plechanovova et al., 2012). While a FANCI-FANCD2-DNA complex is robustly ubiquitinated by Ube2D3 – RNF4^RING-RING^, the Ile44Ala Ub mutant dramatically reduces this modification (Fig 1D). Remarkably, the ubiquitin mutant barely impacts the activity or site-specificity of the Ube2T-FANCL^UR^ pair. These data reveal that while Ube2T–FANCL can catalyse specific ubiquitination it does not share features of the generic RING E3 based catalysis and instead operates through an as yet uncharacterised mechanism.

**Figure 1.**
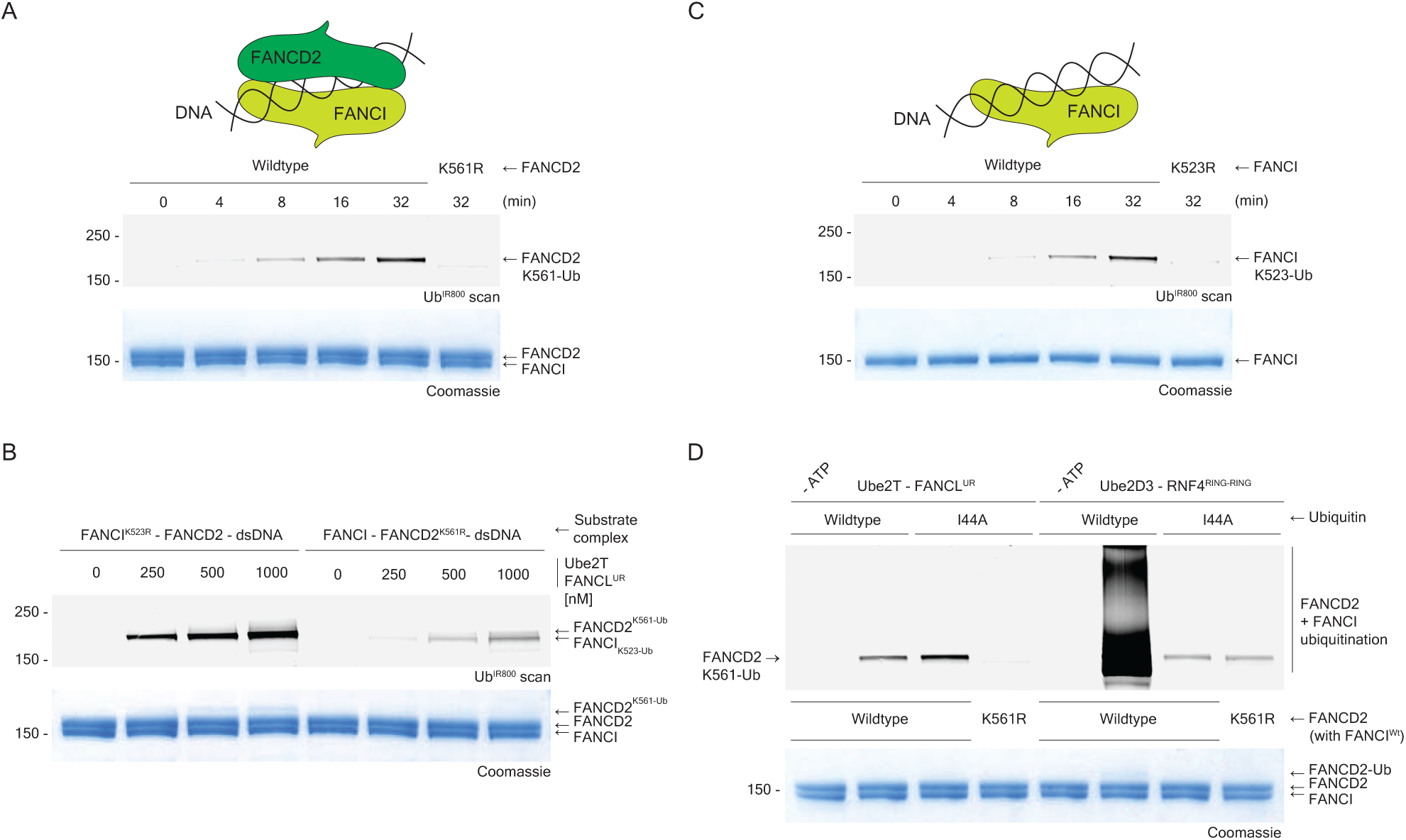
FANCL^UR^ mediated FANCD2 ubiquitination does not require the ubiquitin Ile44-patch. A Time-course multi-turnover ubiquitination assays with fluorescently labelled ubiquitin (Ub^IR800^) showing FANCL^UR^ (0.1 μM) and Ube2T (0.1 μM) mediated site-specific mono-ubiquitination of Lys561 FANCD2 (1.0 μM) when present as a FANCD2-FANCI-dsDNA (1:1:2 μM) complex. B Titration of FANCL^UR^ - Ube2T enzymes on FANCD2-FANCI-dsDNA (1:1:2 μM) complexes with single target lysine i.e. Lys561 FANCD2 or Lys523 FANCI reveals FANCI site can be targeted but FANCD2 site is preferred. C Time-course multi-turnover ubiquitination assays with fluorescently labelled ubiquitin (Ub^IR800^) showing FANCL^UR^ (0.1 μM) and Ube2T (0.1 μM) mediated site-specific mono-ubiquitination of Lys523 FANCI (1.0 μM) when present as a FANCI-dsDNA (1:2 μM) complex. D Comparing FANCD2-FANCI-dsDNA (1:1:2 μM) ubiquitination activities of the FANCL^UR^ (0.1 μM) - Ube2T (0.1 μM) pair with RNF4^RING-RING^ (0.1 μM) - Ube2D3 (0.2 μM) pair using fluorescently labelled wildtype and Ile44Ala ubiquitin. RNF4^RING-RING^ and Ube2D3 robustly ubiquitinates FANCI-FANCD2 using wiltype ubiquitin, but is dramatically impaired by the Ile44Ala mutant while the FANCL^UR^ and Ube2T maintain activity and site-specificity even with the ubiquitin mutant. Substrate ubiquitination is analysed by direct fluorescence monitoring (Li-COR Odyssey CLX).

### FANCL stimulates Ube2T activity through allosteric modulation

As the minimal Ube2T–FANCL^UR^ module drives FANCI and FANCD2 monoubiquitination, we wondered if the underlying mechanism for specific ubiquitination could be uncovered by understanding FANCL’s atypical catalytic mechanism. To investigate this we compared available structures of unbound and the FANCL bound E2. High resolution crystal structures of Ube2T alone (PDB ID 1yh2) (Sheng et al., 2012) and bound to FANCL RING domain (FANCL^R^, PDB ID 4ccg) (Hodson et al., 2014) show little overall difference in their UBC folds (residues 1–152). The latter contains two E2 copies in the asymmetric unit, both of which superpose well onto the unbound state (RMS deviation 0.6–0.9Ǻ). However, RING binding induces some local changes in Ube2T’s helix1–loop2 region as well as in loops 7 and 8 that flank the active site (Fig 2A). Residues in these shifted regions (R3, L7, D32, D33, K91-K95 and D122) are predominantly surface exposed and largely conserved among the Ube2T homologs (Supplementary Fig 1C). We wondered if these subtle conformational changes are important for FANCL’s catalytic mechanism. To test this we made and assayed Ube2T mutants in FANCI and FANCD2 ubiquitination assays. Surprisingly, alanine substitutions of residues Arg3, Asp32/Asp33 and Leu92 reduce the rate of monoubiquitination by 25 to 50% while a loop7 deletion (∆92–95) causes a more severe defect (Fig 2B-C and Supplementary Fig2A). Interestingly, a loop2-loop7 hybrid mutant (D32A, D33A, L92A or DDL/AAA) attenuates both FANCI and FANCD2 ubiquitination rates by around 75%. It is possible that these substrate ubiquitination defects arise from the mutations impairing intrinsic E2 features, such as ubiquitin charging and discharging. The levels of E1-based Ub charging for the above Ube2T mutants is similar to the wildtype E2 (Supplementary Fig 2B). Ube2T readily autoubiquitinates at Lys91, close to the active site (Fig 2A), and several lysines in its C-terminal extension (Machida et al., 2006). We therefore purified an E2 truncation (Ube2T^1–152^) lacking the C-terminal tail. This truncated enzyme targets Lys91 and can be used to assess E2~Ub discharge. In E3-independent Ube2T^1–152^ autoubiquitination assays, a Lys91Arg mutation (Ube2T^1–152^ K91R) abolishes the automodification while the DDL/AAA mutation (Ube2T^1–152^ DDL/AAA) has no observable effect (Fig 2D). Thus in the absence of FANCL, the Ube2T^DDL/AAA^ mutant can load and offload ubiquitin comparable to wildtype Ube2T. Conversely, in single turnover reactions, the mutant E2~Ub thioester (Ube2T^1–152, K91R^ DDL/AAA ~ Ub) is less effective in FANCL^UR^ mediated FANCD2 ubiquitination even in the presence of increasing amounts of the E3 (Fig 2E).

**Figure 2.**
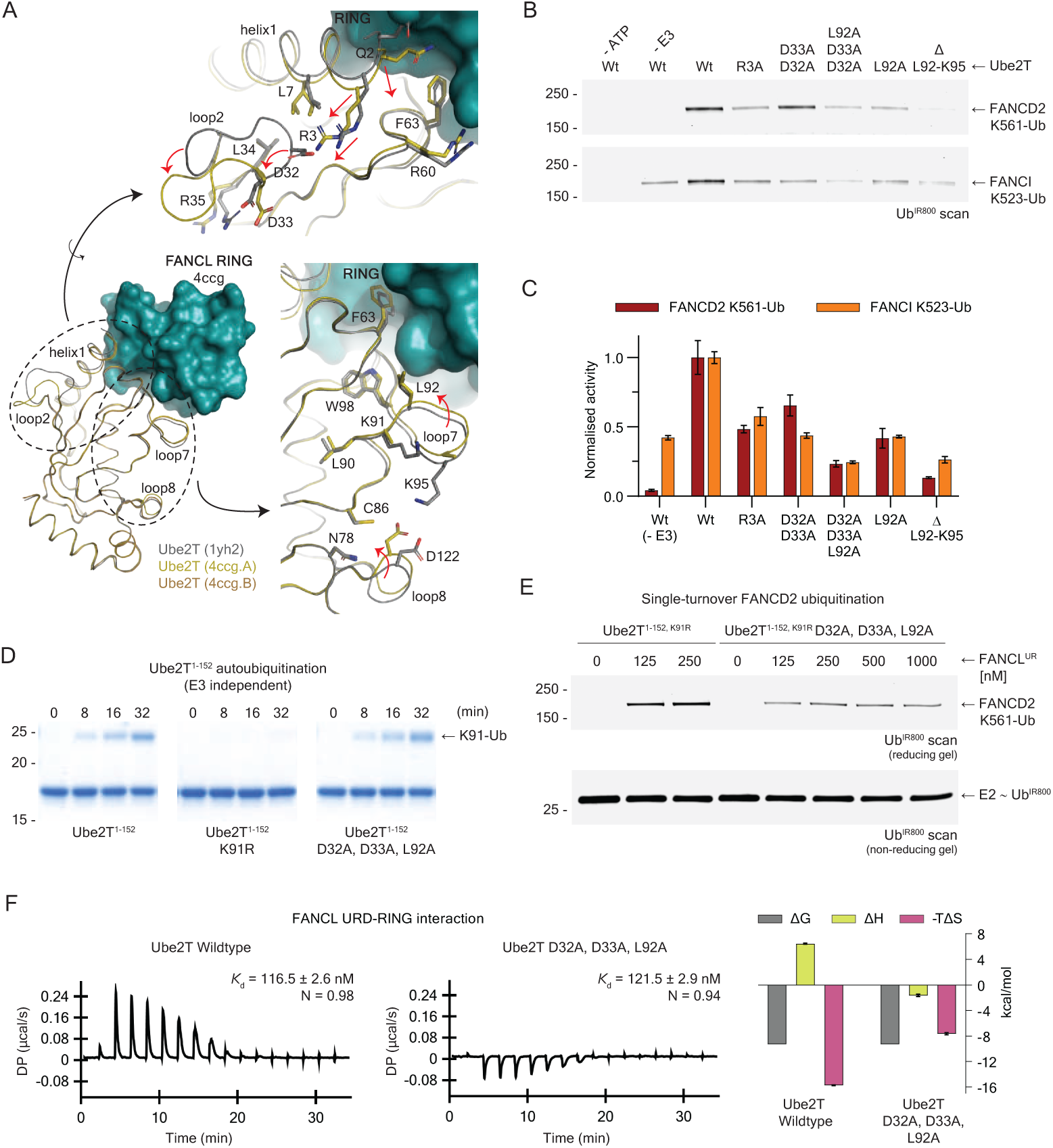
FANCL induced dynamics of Ube2T loop2 and loop7 is required for substrate ubiquitination. A Superpose of FANCL^R^ (teal surface) bound copies of Ube2T (olive ribbon, PDB ID 4ccg.A and brown ribbon, PDB ID 4ccg.B) with the unbound Ube2T (grey ribbon) structure (PDB ID 1yh2) showing little overall structural change. Close-up of helix1-loop2 region (top) and loop7-loop8 region (bottom) reveal local changes. Molecular figures prepared in PyMOL (Schrödinger, LLC). B End-point (30 min) multi-turnover ubiquitination assay with fluorescently labelled ubiquitin (Ub^IR800^) show conserved residues in Ube2T helix1, loop2 and loop7 are required for FANCL^UR^ mediated FANCD2 and FANCI ubiquitination. Substrate ubiquitination is analysed by direct fluorescence monitoring (Li-COR Odyssey CLX). C Effect of Ube2T helix1, loop2 and loop7 mutants on rates of FANCD2 and FANCI ubiquitination. A loop2-loop7 hybrid mutant (Asp32Ala, Asp33Ala, Leu92Ala) shows 75% loss in substrate ubiquitination rates. Rates normalized to wildtype levels and plotted as mean ± SD (n=3). D FANCL independent Ube2T^1–152^ autoubiquitination assay shows no effect of the loop2-loop7 hybrid mutant in Lys91 autoubiquitination. E Single-turnover ubiquitination assay (10 min) of a FANCD2-FANCI^K523R^-dsDNA (2:2:2 μM) complex with increasing amounts of FANCL^UR^ and 200 nM of Ube2T^1–152, K91R^ ~ Ub ^IR800^ thioester or Ube2T^1–152, K91R^ loop2-loop7 hybrid mutant ~ Ub ^IR800^ thioester shows the latter is defective in modifying Lys561 FANCD2. F Thermodynamics of FANCL^UR^ interaction with Ube2T wildtype (left) and Ube2T loop2-loop7 hybrid mutant (middle) shows no change in binding affinity but divergent binding enthalpy. The cost of favourable enthalpy for the mutant is offset by reduced conformational entropy thus contributing to reduced activity. Graphs (right) plotted as mean ± range (n=2).

We wondered if this disparity arises from FANCL being sensed differently by the mutant Ube2T. To uncover possible differences we analysed interactions of wildtype and DDL/AAA mutant Ube2T with the FANCL^UR^ fragment in solution and observe similar affinities (*K*_d_ ~119 nM) indicating that the RING-Ube2T crystal interface is maintained in case of the Ube2T^DDL/AAA^ (Fig 2F). However, the thermodynamics are fundamentally different as the mutant interaction is enthalpically favoured (ΔH = −1.65 kcal/mol) in contrast to the unfavourable signature observed for wildtype Ube2T (ΔH = + 6.46 kcal/mol) (Fig 2F, Table 1). The shorter FANCL^R^ fragment also binds with ~250 nM affinity but shows a similar divergent enthalpy profile (Supplementary Fig 2C). In both FANCL^UR^ and FANCL^R^ experimental sets there appears to be strong enthalpy – entropy compensation and consequently the binding energy (ΔG) within each set is unchanged (Table 1). Therefore, the net differences in observed entropy between wildtype and DDL/AAA Ube2T (8.12 and 6.73 kcal/mol for FANCL^UR^ and FANCL^R^ complexes respectively) could arise from either reduced solvent reordering or fewer local conformational changes in the mutant E2 upon FANCL binding. In other words, the Ube2T^DDL/AAA^ mutant indeed senses FANCL differently from wildtype Ube2T. The biochemical and thermodynamic data taken together reveal that FANCL induces FANCI and FANCD2 ubiquitination by perturbing the overall resting state of Ube2T and strongly suggests allostery. Moreover, FANCL binding indirectly effects Ube2T’s loop2 and loop7 which are 16 – 20Ǻ apart (Fig 2A) and these regions, operating in synergy, propagate the catalytic influence of the E3.

**Table 1.** Thermodynamic properties FANCL - Ube2T interaction^1^. ^1^Average values from two experiments

| E3 | E2 | K<sub>d</sub> (nM) | N | ΔH (kcal.mol<sup>-1</sup>) | ΔG (kcal.mol<sup>-1</sup>) | -TΔS (kcal.mol<sup>-1</sup>) |
| --- | --- | --- | --- | --- | --- | --- |
| FANCL<sup>UR</sup> | Wildtype | 116.5 ± 2.6 | 0.98 | 6.46 ± 0.07 | - 9.30 | -15.75 ± 0.05 |
|  | D32A, D33A, L92A | 121.5 ± 2.9 | 0.94 | -1.65 ± 0.16 | - 9.28 | - 7.64 ± 0.17 |
|  | T23R, W25Q | 200.0 ± 2.7 | 1.00 | 3.90 ± 0.70 | - 8.99 | -12.55 ± 0.15 |
|  | E54R | 119.5 ± 2.2 | 0.93 | 5.60 ± 0.50 | - 9.29 | -14.90 ± 0.10 |
| FANCL<sup>R</sup> | Wildtype | 249.5 ± 7.5 | 1.05 | 3.83 ± 0.36 | - 8.86 | - 12.70 ± 0.40 |
|  | D32A, D33A, L92A | 251.5 ± 1.5 | 0.95 | - 2.88 ± 0.02 | - 8.86 | - 5.98 ± 0.03 |
|  | T23R, W25Q | 259.0 ± 8.4 | 1.05 | 3.89 ± 0.05 | - 8.84 | - 12.75 ± 0.05 |

<sup>1</sup>Average values from two experiments

### Ube2T backside regulates FANCL mediated FANCD2 ubiquitination

Our binding analyses reveal a 2-fold enhanced Ube2T affinity for the FANCL^UR^ fragment over the smaller RING domain (Table 1, Supplementary Fig 2C) indicating additional interactions exist between a non-RING element and the E2. In several RING E3s, auxiliary elements outside of the RING domain can modulate ubiquitination by binding an E2 ‘backside’ surface which is located opposite the active site (Brown et al., 2015; Das et al., 2009; Hibbert et al., 2011; Li et al., 2015; Li et al., 2009; Metzger et al., 2013; Turco et al., 2015). We wondered if the analogous Ube2T backside surface could extend the FANCL-E2 interface as well as influence FANCI and FANCD2 ubiquitination. Beta-strands 1 and 2 of the UBC-fold prominently feature in E3 – backside E2 complexes (PDB IDs 3h8k, 2ybf, 4jqu, 5d1k and 4yii) and at the canonical ubiquitin – backside E2 interface (PDB IDs 2fuh and 4v3l). The equivalent Ube2T surface is hydrophobic, semi-conserved (Supplementary Fig 1C) with certain side-chains (*β*1 – T23, W25 and *β*2 – R35, Q37) repositioned upon FANCL^R^ binding (Fig 3A). Notably, the mutation of Ube2T *β*1 (T23R+W25Q or TW/RQ) reduces the E2’s affinity for FANCL^UR^ (*K*_d_ – 200 nM) but not for FANCL^R^ (Fig 3B, Table 1). Thus, the Ube2T backside indeed supports additional interactions with FANCL^UR^ beyond the RING domain. Unexpectedly, the backside mutants have different effects on substrate ubiquitination. Previous *in vitro* studies have reported site-specific ubiquitination of a FANCI/DNA complex by Ube2T in the absence of FANCL (Longerich et al., 2014). In our setup, FANCL^UR^ enhances FANCI ubiquitination rates by around two-fold while backside Ube2T mutants mitigate this improvement (Fig 3C-D and Supplementary Fig 3A). In contrast, the Ube2T TW/RQ mutant slows FANCD2 ubiquitination by over three-fold, similar to the DDL/AAA mutant (Fig 2B-C). We also tested a *β*2 mutant (Q37L), designed to extend the hydrophobic backside surface, and observe little change in ubiquitination rates (Fig 3C, D). Overall, the backside Ube2T mutants do not affect ubiquitin charging or Ube2T^1–152^ auto-ubiquitination (Supplementary Fig 3A and 3E) and therefore indicate the ubiquitin loading/offloading properties of the mutants are intact. Hence, the observed defects in substrate modification are likely linked to an altered FANCL^UR^-backside Ube2T interaction. We wondered if the weakened Ube2T TW/RQ-FANCL^UR^ interaction is solely responsible for reduced FANCD2 ubiquitination. In single-turnover reactions, the charged Ube2T^1–152, K91R^ TW/RQ ~ Ub thioester is weakly activated by FANCL^UR^ for FANCD2 ubiquitination however, increasing the amount of E3 does not completely rescue the defect (Fig 3F). Thus, Ube2T’s backside surface not only supports FANCL^UR^ interaction but likely augments the allosteric activation of the E2~Ub thioester by the E3. In summary, loop2, loop7 and the backside of Ube2T together respond to FANCL binding to facilitate FANCI and FANCD2 ubiquitination.

**Figure 3.**
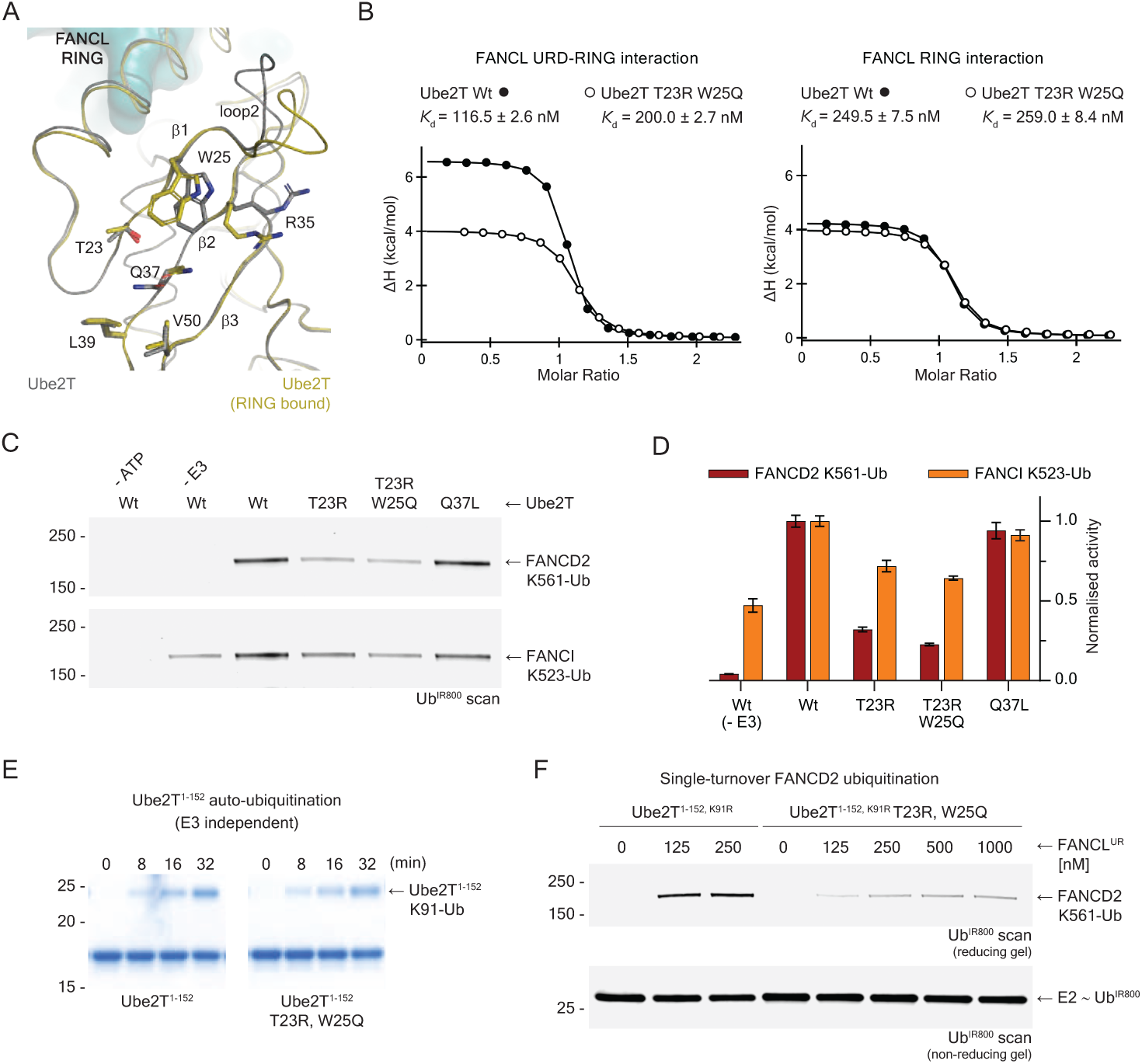
Novel role of Ube2T backside in FANCL mediated substrate ubiquitination. A Superpose of FANCL^R^ (teal surface) bound copy of Ube2T (olive ribbon, PDB ID 4ccg.A) with unbound Ube2T (grey ribbon) structure (PDB ID 1yh2) showing residues on E2 backside (β1 and β2) that are repositioned upon FANCL binding. Molecular figures prepared in PyMOL (Schrödinger, LLC). B Ube2T backside β1 double mutant (Thr23Arg, Trp25Gln) reveals binding defect with FANCL^UR^ but is unaffected in FANCL^R^ interaction. This suggests Ube2T backside supports additional FANCL interactions. C End-point (30 min) multi-turnover ubiquitination assay with fluorescently labelled ubiquitin (Ub^IR800^) show Ube2T backside mutants are more defected in FANCL^UR^ mediated FANCD2 ubiquitination than FANCI ubiquitination. Substrate ubiquitination is analysed by direct fluorescence monitoring (Li-COR Odyssey CLX). D Effect of Ube2T backside mutants on the rates of FANCD2 and FANCI ubiquitination. Backside mutants reduce FANCI modification rate to levels observed in the no E3 setup. In contrast, a Ube2T backside β1 double mutant (Thr23Arg, Trp25Gln) shows 75% loss in FANCD2 ubiquitination. Rates normalized to wildtype levels and plotted as mean ± SD (n=3). E FANCL independent Ube2T^1–152^ autoubiquitination assay shows no effect of the backside β1 double mutant in Lys91 autoubiquitination. F Single-turnover ubiquitination assay (10 min) of a FANCD2-FANCI^K523R^-dsDNA (2:2:2 μM) complex with increasing amounts of FANCL^UR^ and 200 nM of Ube2T^1–152,K91R^ ~ Ub ^IR800^ thioester or Ube2T^1–152, K91R^ β1 mutant ~ Ub ^IR800^ thioester shows the defect in the latter cannot be completely rescued by increasing FANCL^UR^ levels.

### FANCL potentiates Ube2T active site residues for FANCI AND FANCD2 ubiquitination

The above data reveal how the FANCL influence on distinct Ube2T surfaces triggers the E2~Ub for substrate ubiquitination. Thus a long-range residue network could connect these distal sites with the E2 catalytic centre. To uncover the likely path we generated residue interaction networks (RINs) for the free and RING bound Ube2T structures (residues 1–152) (Doncheva et al., 2011; Piovesan et al., 2016). In these networks, each Ube2T residue is represented as a node while the connecting edges are potential physicochemical interactions with its tertiary structure environment. The total connections in both free and RING bound E2 RINs are similar, averaging 1430 edges, however a comparison matrix reveals unchanged and altered edges (Fig 4A, Supplementary Dataset 1), the latter used to build a dynamic network. We then choose Phe63, a core E2 residue at the Ube2T-FANCL^R^ interface (Hodson et al., 2014), as the starting node to trace its first neighbours which serve as subsequent search nodes. By iteration, we trace the possible paths to the E2 catalytic centre, focusing on the allosteric and conserved nodes while filtering out paths comprising distal and dead-end nodes. The final allosteric network model (39 nodes, 79 dynamic edges) reveals how FANCL^R^ binding rewires Ube2T’s intra-molecular connections (Fig 4A, Supplementary Table 1).

**Figure 4.**
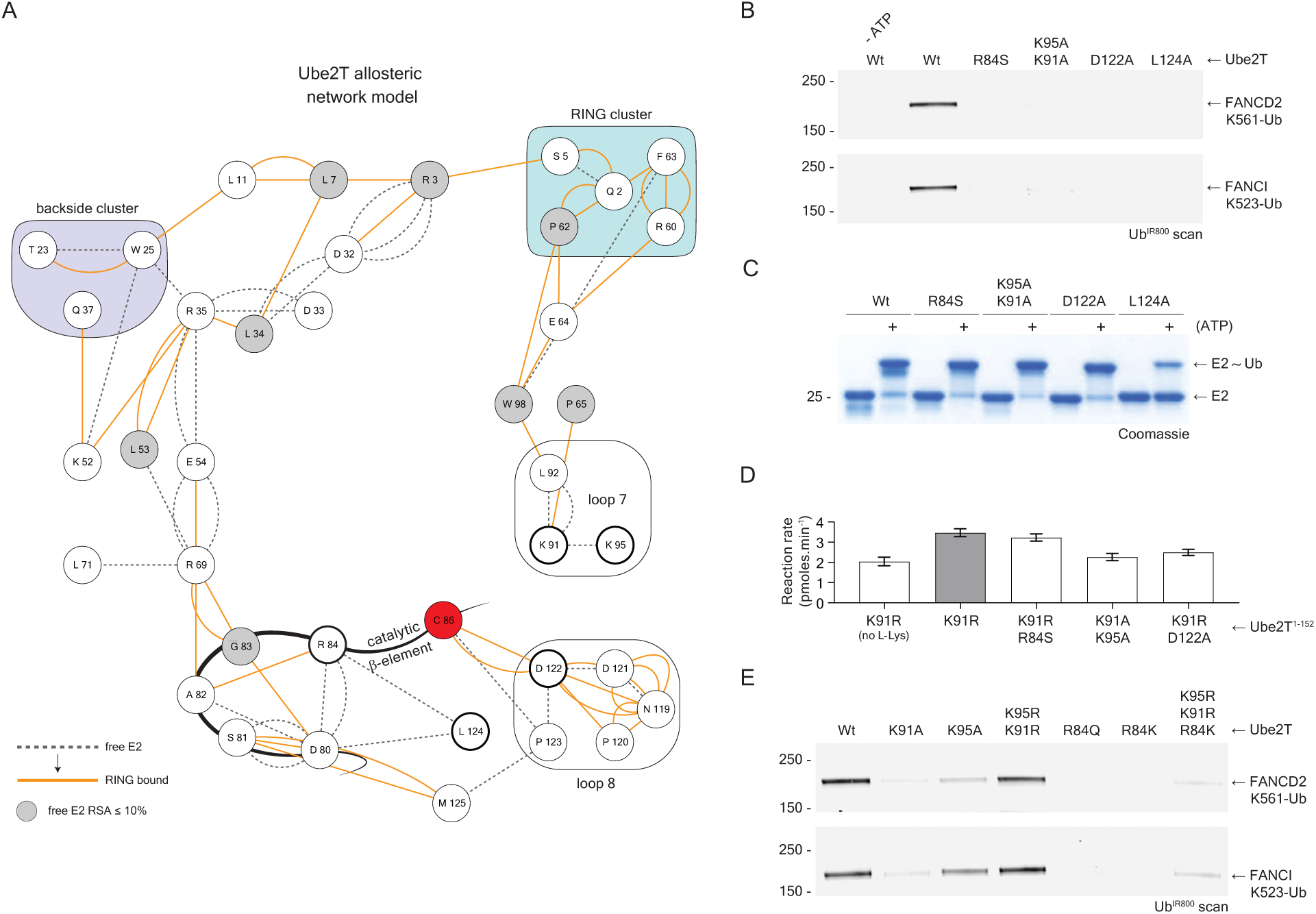
Allosteric residue network reveals Ube2T active-site residues critical for substrate ubiquitination. A Allosteric network model shows dynamic rewiring of Ube2T intra-residue connections upon FANCL^R^ binding. Dashed and orange lines depict Ube2T edges in unbound and FANCL^R^ bound state respectively. Grey nodes have relative solvent accessibility of less than 10% in unbound Ube2T. Nodes involved in RING binding are clustered in a cyan box while those predicted to support backside interaction are clustered in a purple box. Nodes in loop7 and loop8 are in unshaded boxes. Red node denotes the catalytic cysteine (Cys86) while the network termini nodes (Arg84, Lys91, Lys95, Asp122 and Leu124) that are within 10 Ǻ of Cys86 have a thick outline. Also depicted is the catalytic beta-element. B End-point (30 min) multi-turnover ubiquitination assay with fluorescently labelled ubiquitin (Ub^IR800^) show mutations of the Ube2T network termini residues are detrimental to FANCL^UR^ mediated FANCD2 and FANCI ubiquitination. Substrate ubiquitination is analysed by direct fluorescence monitoring (Li-COR Odyssey CLX). C Ubiquitin charging assays of the network termini mutants show the Leu124Ala mutant alone is defected E1-based E2~Ub thioester formation. D Lysine discharge assays show network termini mutants in loop7 (Lys91Ala + Lys95Ala) and loop8 (Asp122Ala) have catalytic defects while Arg84Ala is not defected. Graphs depict mean ± SD (n=2). E End-point (30 min) multi-turnover ubiquitination assay with fluorescently labelled ubiquitin (Ub^IR800^) show the requirement of Arg/Lys residues in loop7 while Arg84, in the catalytic β-element, is critical for FANCI and FANCD2 ubiquitination. Partial compensation of activity for Ube2T Arg84Lys occurs only when the loop7 bears longer Arg residues.

The network terminals, located in the catalytic beta-element (R84), loop7 (K91, K95), loop8 and its C-terminal hinge (D122 and L124 respectively) are within 10Ǻ of Ube2T’s catalytic cysteine (C86) and vary among the ubiquitin E2s (Fig 4A and Supplementary Fig 4B). To empirically test the network model we made alanine mutants of the said network termini and observe a striking loss in FANCL mediated FANCI and FANCD2 ubiquitination (Fig 4B). Given their proximity to the active site these Ube2T residues could also influence intrinsic E2 activity. The Leu124Ala mutation is detrimental to E1-based ubiquitin charging and suggests the hydrophobic side-chain braces Ube2T’s active site for optimal activity (Fig 4C). Moreover, by examining E3-independent E2~Ub thioester discharge onto free lysine, we observe subtle catalytic defects with the loop7 (Ube2T^1–152^ K91A+K95A) and loop8 (Ube2T^1–152, K91R^ D122A) mutants while in contrast, the Arg84Ser (Ube2T^1–152, K91R^ R84S) mutant did not affect Ube2T’s aminolysis activity (Fig 4D and Fig Supplementary Fig 4C). In some E2s, the loop8 acidic residue is required to position and/or deprotonate the target lysine for modification (Plechanovova et al., 2012; Valimberti et al., 2015; Yunus and Lima, 2006). The loop8 Asp122 in Ube2T could support a similar role and account for the catalytic defects with this mutant. However, the contrasting E2 activity profiles (E3-independent free lysine versus E3-dependent substrate lysine, Fig 4B and D) for the catalytic beta-element mutant (R84S) suggests the Arg84 residue is likely involved in FANCL’s activation mechanism.

Moreover, as we earlier observe the Leu92Ala loop7 mutant of Ube2T reduces FANCL mediated substrate ubiquitination (Fig 2B-C) we reasoned the E3’s activation mechanism feeds into the catalytic role of this loop (Fig 4A, D).

To understand roles of Ube2T residues Arg84, Lys91 and Lys95 in substrate ubiquitination we undertook systematic mutagenesis and uncover the requirements of Lys/Arg in loop7 while the Arg84 residue is indispensable for FANCL mediated FANCI and FANCD2 ubiquitination (Fig 4E). To clarify this necessity we analysed the total network (unchanged and altered) for the Arg84 node (Supplementary Fig 4D). In free Ube2T, the Arg84 side-chain stabilises the Asp80-Gly83 loop and is transiently redirected towards Cys86 upon RING binding (Fig 4A, Supplementary Fig 4B and 4D). A Arg84Lys mutation would, in theory, preserve the free E2 network but the persistent defect in substrate ubiquitination (Fig 4E) suggests the shorter Lys side-chain is unable to optimally participate at the active site. Interestingly, the introduction of longer Arg side-chains in loop7 (R84K, K91R, K95R) can rescue the Arg84Lys defect, albeit partially (Fig 4E). In summary, FANCL binding potentiates the Ube2T active site residues R84, K91 and K95 to induce FANCI and FANCD2 ubiquitination. These residues could enhance catalysis either through transient interactions with acidic/polar substrate residues proximal (<12Ǻ) to the respective target lysine (Supplementary Fig 4C) and/or stabilise the developing oxyanion hole in the E2~Ub – target lysine transition state. Taken together, the data reveals the existence of an elaborate Ube2T residue network that propagates FANCL’s catalytic influence in activating the E2~Ub thioester for ubiquitination.

### Ube2T network analysis reveals regulatory and FANCL induced activation mechanisms

The thermodynamics of Ube2T-FANCL interaction and the allosteric network model together illustrate how the E3 could remotely effect the E2 catalytic centre by inducing a series of subtle conformational changes within Ube2T. Intriguingly, we noticed a dynamic conduit involving nodes Arg35 – Glu54 – Arg69 that appear to propagate FANCL’s influence on the RING and backside clusters to the catalytic beta-element (Fig 4A, Supplementary Table 1). In particular, the Glu54 side-chain which engages both Arg35 and Arg69 in unbound Ube2T has a reduced influence in the RING bound state. Subsequently, the Arg69 side-chain is liberated to stabilise the catalytic beta-element backbone thus releasing the Arg84 side-chain to optimally participate at the catalytic centre (Fig 5A). In other words, an Arg69 effector role in Ube2T’s resting state could be gated by Glu54 and relieved upon FANCL binding thereby activating the E2~Ub for FANCI and FANCD2 ubiquitination. Based on this model we predict the removal of residue 54’s acidic side-chain should positively impact substrate ubiquitination. We tested the proposed allosteric conduit using Glu54Ala/Gln mutations and observe a marked improvement in FANCL^UR^ mediated FANCI AND FANCD2 ubiquitination while a conservative Glu54Asp mutant retained wild-type like activity (Fig 5B). Even a charge-altering Glu54Arg mutation contributes to greater substrate ubiquitination and does not influence the Ube2T-FANCL^UR^ interaction (Fig 5B, Table 1). Consequently, an Arg69Ala mutation reduces FANCI and FANCD2 ubiquitination, while a conservative mutation (R69K) rescues this defect and is further improved by eliminating the gating effect (E54A with R69K) thus confirming residue 69’s effector role (Fig 5C). Furthermore, disrupting the Glu54 gate can rescue FANCD2 ubiquitination defects arising from both RING allostery (DDL/AAA) and backside binding (TW/RQ) mutants (Fig 5D). Interestingly, a RING binding E2 mutant (F63A) that reduces Ube2T-FANCL^R^ binding by over ten-fold (Hodson et al., 2011) could also be activated by FANCL^UR^ for FANCD2 ubiquitination by using a permissive gate (F63A with E54R) (Fig 5D).

**Figure 5.**
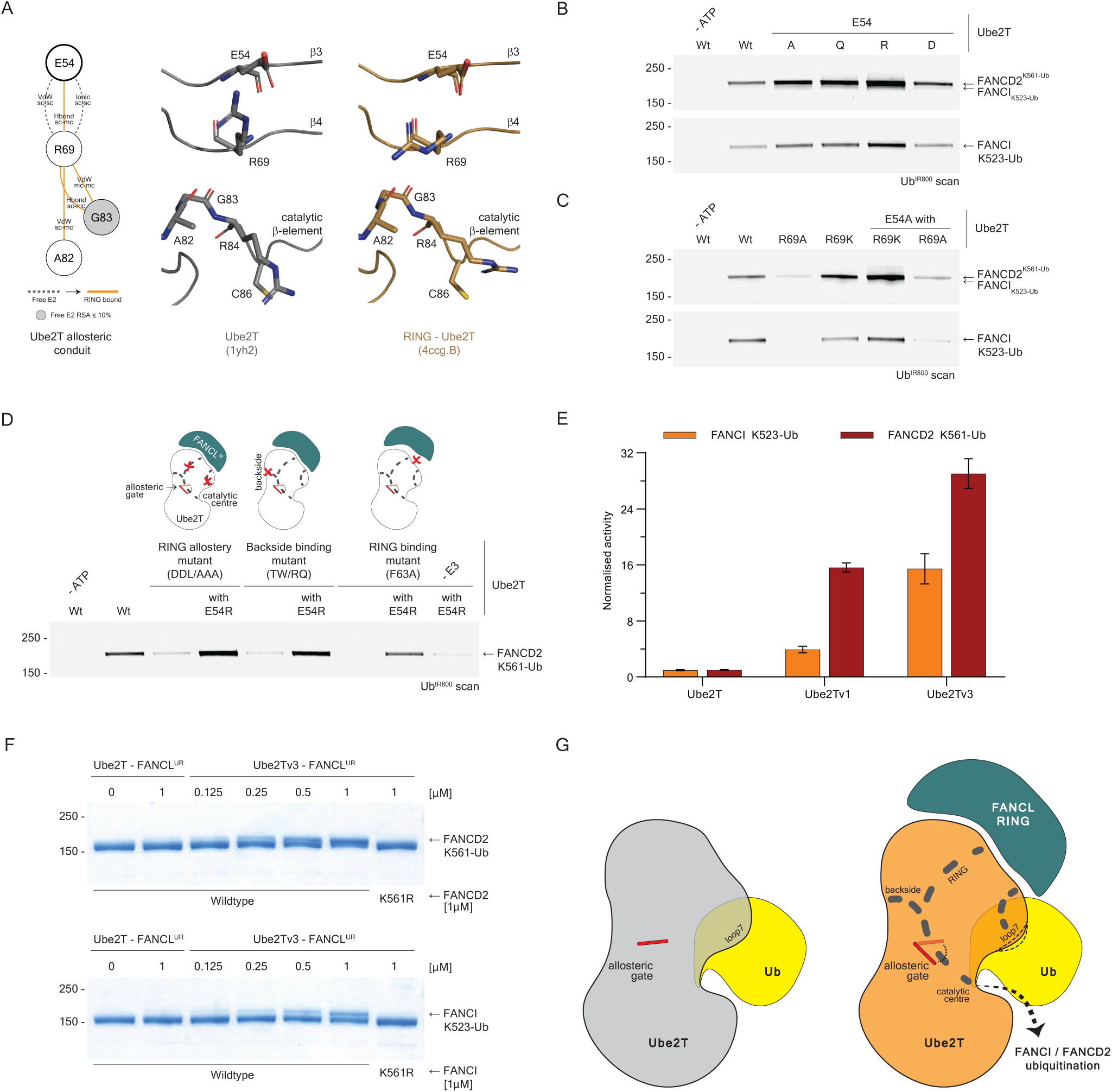
Ube2T deregulation leads to enhanced FANCL driven substrate ubiquitination. A Comparison of a network scheme (left) with structures of unbound (middle) and RING bound (right) Ube2T depicting the proposed allosteric conduit. A gating role is proposed for Glu54 (thick outline) for its regulation of Arg69. An effector role is proposed for Arg69 for its role in stabilising the catalytic β-element leading to a release of Arg84. Dashed and orange lines depict Ube2T edges in unbound and FANCL^R^ bound state respectively. Grey nodes have relative solvent accessibility of less than 10% in unbound Ube2T. Molecular figures prepared in PyMOL (Schrödinger, LLC). B End-point (30 min) multi-turnover ubiquitination assay with fluorescently labelled ubiquitin (Ub^IR800^) shows Glu54 gating role is dependent on its negative side chain. Removal of this charge leads to improved FANCL^UR^ mediated substrate ubiquitination. Substrate ubiquitination is analysed by direct fluorescence monitoring (Li-COR Odyssey CLX). C End-point (30 min) multi-turnover ubiquitination assay with fluorescently labelled ubiquitin (Ub^IR800^) shows effector role for Arg69 is linked to its positive side chain and is regulated by the Glu54 gate. While removal of the Glu54 negative charge improves improved FANCL^UR^ mediated substrate ubiquitination, the removal of the Arg69 positive charge counters this effect. D End-point (30 min) multi-turnover ubiquitination assay of a FANCD2-FANCI^K523R^- dsDNA (1:1:2 μM) complex with fluorescently labelled ubiquitin (Ub^IR800^) shows a permissive gate at position 54 (Glu54Arg) can rescue FANCD2 ubiquitination defects arising from the loop2-loop7 hybrid mutant (Asp32Ala, Asp33Ala, Leu92Ala), the backside β1 double mutant (Thr23Arg, Trp25Gln) as well as the RING binding mutant (Phe63Ala). Despite deregulation of the RING binding Ube2T Glu54Arg + Phe63Ala double mutant, FANCL is still required for efficient FANCD2 ubiquitination. E Effects of the Ube2Tv1 variant with a permissive gate (Glu54Arg) alongside the Ube2Tv3 variant containing a permissive gate and flexible loop7 (Glu54Arg, Pro93Gly, Pro94Gly) on the rates of FANCL^UR^ mediated FANCD2 (FANCI^K523R^- FANCD2-dsDNA complex) and FANCI (FANCI-dsDNA complex) ubiquitination. F End-point (30 min) multi-turnover ubiquitination assay of isolated FANCD2 and FANCI substrates with no DNA cofactors. Titration of FANCL^UR^ - Ube2Tv3 enzymes shows how substrate ubiquitination is enhanced by the deregulated E2 without compromising on site-specificity. G Allosteric model for FANCL^UR^ mediated activation of Ube2T. FANCL binding at classical RING-E2 interface and the backside E2 interface triggers long-range and short-range rewiring of Ube2T residue networks that culminate in substrate ubiquitination. The dynamic allosteric network is intrinsically regulated by conserved Ube2T elements, the allosteric β3 gating residue and a rigid loop7.

We wondered if such gated networks exists in other E2s, in particular those that assemble specific ubiquitination signals. For example, DNA damage tolerance pathways are initiated upon specific Lys164 monoubiquitination of Proliferating Cell Nuclear Antigen (PCNA) protein by the E2-E3 enzyme pair Ube2B/Rad6-RAD18 (Ulrich and Walden, 2010). The RAD18-Ube2B interaction is bimodal, primarily via the canonical RING-E2 interface and supported by a helical Rad6 binding domain (R6BD) that packs against the E2’s backside (Hibbert et al., 2011; Huang et al., 2011). We compared the R6BD bound (PDB ID 2ybf) structure/RIN with the free Ube2B (PDB ID 2yb6) and noticed Glu58 and Arg71 of the E2 could potentially operate as gating and effector residues respectively (Supplementary Fig 5A, Supplementary Dataset 2). Notably in *in vitro* assays, the Ube2B variant with a permissive gate (E58R) is more sensitive to Rad18 in PCNA monoubiquitination without comprising site-specificity (Supplementary Fig 5A). These results together demonstrate that different E2’s contain long-range RINs that are regulated by internal gates. Furthermore, these networks can be leveraged by RING E3 ligases to allosterically drive ubiquitination of specific targets.

A short-range RIN is also apparent between Ube2T residues that support RING binding and those in loop7 (Fig 4A, Supplementary Table 1). The binding of FANCL^R^ draws loop7 (K_91_LPPK_95_) away from the active site (Fig 2A) thereby repositioning the flanking Lys residues (K91 and K95) presumably for target site binding, while paradoxically these residues are also needed at the active site for optimal Ube2T catalysis (Fig 4C). Also present in this loop are two conserved proline residues (P93 and P94, Supplementary Fig 1C) that maintain loop rigidity and we hypothesise that a plastic loop7 could emulate its dynamic requirements during ubiquitination. In multi-turnover assays, the Ube2Tv1 variant with a permissive gate (E54R) improves FANCL^UR^ driven FANCD2 ubiquitination by around 15- fold (around 4-fold for FANCI) while a hybrid Ube2Tv3 variant that includes both a permissive gate and flexible loop7 (E54R, P93G, P94G) further enhances activity by 2 to 4- fold (Fig 5E and Supplementary Fig5B) thus validating our hypothesis. The above data reveal distinct regulatory mechanisms in operation within Ube2T’s allosteric network and these can be repurposed to yield E2 variants with enhanced catalytic potential. Furthermore, the enhanced Ube2Tv3 – FANCL^UR^ pair facilitates steady and site-specific ubiquitination of the isolated FANCD2 and FANCI proteins in the absence of any DNA cofactors (Fig 5F). Recent reports reveal the *in-vitro* requirements of a large (~ 0.8 MDa), multi-protein E3 super-assembly (FANCB-FANCL-FAAP100-FANCC-FANCE-FANCF) that activates the Ube2T~Ub thioester for robust ubiquitination of a FANCI-FANCD2-DNA complex (Swuec et al., 2017; van Twest et al., 2017). By investigating the catalytic mechanism of the core E2 – E3 proteins within this super-assembly i.e. Ube2T – FANCL, we have rationally engineered a minimal module (Ube2Tv3 – FANCL^UR^, ~ 0.05 MDa) with enhanced catalytic ability to autonomously, specifically and efficiently mono-ubiquitinate FANCI or FANCD2. Using this simplified setup we can now systematically examine roles of the FA core-complex members in ubiquitination, biochemically and structurally characterize the natively ubiquitinated FANCI and FANCD2 substrates, identify novel readers of the ubiquitin signal that implement DNA repair as well as understand how this specific signal is removed by the cognate deubiquitinating enzyme USP1 (Nijman et al., 2005).

## Discussion

Understanding the mechanisms for site-specific FANCD2 and FANCI mono-ubiquitination would shed light on how this decisive DNA damage response signal is mediated by the multi-protein FA core-complex as well as divulge ubiquitination strategies used in precision targeting. Our previous studies on FANCL, the catalytic RING-bearing subunit in the FA core-complex, revealed substrate binding via its central UBC-RWD domain (Hodson et al., 2011). Moreover, an extended RING-E2 interface underlies the strong E3-E2 affinity and enables FANCL to specify its E2 partner Ube2T (Hodson et al., 2014). However, the mechanism by which FANCL catalyses monoubiquitination at specific FANCD2 and FANCI sites, a key signal in the FA-ICL repair pathway (Ceccaldi et al., 2016), is poorly understood. In this study we expand the functional significance of the intimate FANCL-Ube2T interaction and uncover an atypical mechanism behind substrate ubiquitination.

Unlike other RING E3s, FANCL lacks the conserved linchpin residue (Arg/Lys) essential for stabilizing a closed and productive E2~Ub conformation (Metzger et al., 2014). This suggests FANCL’s RING domain may not activate most E2~Ub intermediates thus precluding any non-specific modification or the assembly polyubiquitin signals on the FANCI and FANCD2 substrates. Despite missing this feature FANCL does activate Ube2T for site-specific substrate ubiquitination. Using thermodynamic and residue network analysis we demonstrate how FANCL’s high-affinity grasp of Ube2T induces a series of subtle conformational changes within the E2 that are relevant for substrate ubiquitination. These changes transpire through altered intra-residue connections between conserved Ube2T residues, revealing a dynamic allosteric network that links FANCL binding to the E2 catalytic centre. Notably, conserved basic residues (Arg84, Lys91 and Lys95) proximal to Ube2T’s catalytic cysteine (Cys86) are repositioned in the FANCL^R^ bound structure. While these residues partially contribute to intrinsic Ube2T activity, they are critical for FANCL induced FANCD2 and FANCI ubiquitination. Several studies on site-specific histone monoubiquitination mechanisms show how interactions between RING domain residues and the substrate surface guide the catalytic RING-E2~Ub complex to a lysine targeting zone (Bentley et al., 2011; Gallego et al., 2016; Mattiroli et al., 2014; Mattiroli et al., 2012; McGinty et al., 2014). The FANCL-Ube2T complex is similar in that substrate docking by FANCL’s UBC-RWD module could restrict the global targeting area of the RING bound Ube2T. However, unique to this E3-E2 pair is FANCL RING allostery which reorients the basic residues near Ube2T’s active-site, which could facilitate local contacts with conserved acidic/polar FANCD2/FANCI residues in the vicinity of the target lysine, thus directing specific ubiquitination. It remains to be seen if other RING E3s can actively repurpose E2 residues for specific lysine targeting. However, this strategy of E2-guided lysine targeting mirrors those observed in autonomous polyubiquitin assembling E2s where the acceptor ubiquitin surface around the target lysine is homed by auxiliary E2 interactions (Eddins et al., 2006; Liu et al., 2014; Middleton and Day, 2015; Petroski and Deshaies, 2005; Rodrigo-Brenni et al., 2010; Wickliffe et al., 2011). Alternatively, given the necessity of DNA cofactors in FANCI/FANCD2 modification (Liang et al., 2016; Longerich et al., 2014; Sato et al., 2012), a DNA sensing role for these Ube2T basic residues cannot be ruled out. In either scenario, post-ubiquitination the local target zone for the FANCL-Ube2T complex is occluded by the attached mono-ubiquitin. As we do not observe continual modification, either on a different substrate lysine or the installed ubiquitin, we propose neither surface is efficiently recognized by the FANCL-Ube2T complex thus limiting the ubiquitination to a single event.

Further, we uncover residues on Ube2T’s backside (β1 and β2) that support additional interactions with FANCL and feature in the E2’s allosteric network. It is likely that the backside Ube2T binding is mediated by FANCL’s UBC-RWD domain which also docks onto the substrate surface. As the backside interaction is required for steady substrate ubiquitination, FANCD2 in particular, we speculate that additional UBC-RWD and Ube2T interactions could guide the E2~Ub to the substrate as well as allosterically activate the enzyme. Notably, these observations enlist Ube2T to the growing number of E2’s whose ubiquitination activities are modulated by interactions outside of the classical RING-E2 interface (Brown et al., 2015; Das et al., 2009; Hibbert et al., 2011; Li et al., 2015; Li et al., 2009; Metzger et al., 2013; Turco et al., 2015). Finally, the network guided biochemical analysis also reveals the presence of dynamic long-range and short-range residue networks that are intrinsically regulated by conserved Ube2T residues. In particular, an allosteric gating residue in β3 and a rigid loop7 appear to regulate the intensity of Ube2T activation by FANCL. Rationally engineered mutations of these regions give rise to deregulated Ube2T variants which are strikingly more responsive to FANCL in substrate ubiquitination, yet retain the lysine specificity observed with the wildtype E2. Notably, the catalytic enhanced Ube2T variants now support FANCL driven monoubiquitination of the isolated FANCD2 and FANCI proteins without needing DNA cofactors. These data reveal the molecular strategies in place in Ube2T that prevent inadvertent and untimely mono-ubiquitin signals in the FA-ICL repair pathway. Taken together, we propose a model where FANCL binding at the canonical RING-E2 interface and the backside E2 surface rewires Ube2T’s residue network and thus activates the enzyme for site-specific FANCI and FANCD2 monoubiquitination (Fig. 5G).

On the basis of Ube2T’s allosteric network we identify a similar regulated network in Ube2B/Rad6 that is altered by the RING E3 Rad18 for Lys164 PCNA mono-ubiquitination. Biochemical, structural and computational studies have revealed how RING/U-box E3s stimulate internal dynamics in different E2s that are linked to their ubiquitination activity (Benirschke et al., 2010; Chakrabarti et al., 2017; Das et al., 2013; Ozkan et al., 2005). Furthermore, recent efforts in fragment-based inhibitor discovery have revealed promising lead compounds that bind Ube2T (Morreale et al., 2017) as well as the unrelated Ube2I (Hewitt et al., 2016) at regions distal from their active/E3 binding sites, nevertheless can allosterically regulate the respective E2 activities. These observations collectively suggest that allosteric networks could operate across the E2 family and that future investigations into such networks could prove instrumental for basic and translational research in ubiquitin biology. Using our Ube2T-gated network as a guide and the available E2 sequence/structural data, we propose at least 8 other ubiquitin E2s (Ube2- A, C, E1, E2, E3, K, L3 and N) contain a β3 gating residue that is restrictive (Supplementary Fig 5C). In contrast, the small ubiquitin-like modifier (SUMO) E2 Ube2I and Interferon-stimulated gene 15 (ISG15) E2 Ube2L6 appear to contain a permissive β3 gate. Interestingly, opposing allosteric gates are apparent for the two neural precursor cell expressed developmentally downregulated protein 8 (NEDD8) E2s Ube2F (restrictive) and Ube2M (permissive). Since the β3 gating residue does not lie in the typical RING or backside binding regions, we speculate that targeting this residue would maintain E2-E3 interactions nevertheless, will yield E2 variants that differ in their enzymatic potential relative to their wildtype counterparts. Recent studies have utilized phage-display derived ubiquitin binding variants to uncover mechanistic insights for RING/U-box (Gabrielsen et al., 2017) and HECT (Zhang et al., 2016) E3s as well as deubiquitinating enzymes (Ernst et al., 2013). Along these lines, we propose the design principles for creating E2 activity variants that can be used in fundamental research and guide drug discovery studies.

## Materials and Methods

### Cloning and mutagenesis of expression constructs

Human Ube2T and Ube2D3 was cloned using PCR/restriction cloning into a modified pET15 vector (Novagen) that express with a 6xHis-3C cleavage site at the N-terminus. Human Ube2B cDNA was as an I.M.A.G.E. clone (Geneservice) and cloned into a modified pDEST17 vector (Invitrogen), that express with a 6xHis-TEV cleavage site at the N-terminus using the Gateway Cloning kit (Invitrogen). A synthetic human FANCL sequence (GeneArt) optimized for *Escherichia coli* (*E. coli*) expression used as template to clone the FANCL^UR^ (residues 109–375) and FANCL^R^ (residues 289–375) coding regions into a pET SUMO (Invitrogen) expression vector using restriction-free cloning (van den Ent and Lowe, 2006). These constructs express with a 6xHis-smt3 tag at the N-terminus. A synthetic human Rad18 sequence (GeneArt) optimized for *Escherichia coli* expression was cloned using PCR/restriction cloning into a modified pET28a (Novagen), that express with 6xHis-smt3 tag at the N-terminus. A synthetic Human FANCD2 sequence (GeneArt) optimized for *Spodoptera frugiperda* (*Sf*) expression was cloned using PCR/restriction cloning into a pFastBac vector that express with a 6xHis-3C cleavage site at the N-terminus. Human FANCI (cDNA purchased as an I.M.A.G.E. clone, Geneservice) was cloned using PCR/restriction cloning into a pFastBac vector that express with a 6xHis-TEV cleavage site-V5 epitope at the N-terminus. Human PCNA template, a kind gift from Dr Svend Petersen-Mahrt (European Institute of Oncology, Milan), was cloned using PCR/restriction cloning into a pRSF Duet1 vector (Novagen), that express with a 6xHis at the N-terminus. Ubiquitin was cloned using PCR/restriction cloning into a modified pET28a vector (Novagen), that express with 6xHis-smt3 tag-SGCGSG overhang at the N-terminus for fluorescence labelling or into a modified pRSF Duet1 vector (Novagen), that express with a 6xHis-TEV cleavage site at the N-terminus. All desired mutagenesis were carried out by primer based PCR mutagenesis using KOD Hot Start DNA polymerase (Novagen) or Phusion High-Fidelity DNA polymerase (Thermo Scientific) following manufacturer’s guidelines. Custom oligonucleotides for PCR and mutagenesis were obtained from Sigma Aldrich or IDT technologies. DH5α *E. coli* strain were made chemically competent using CCMB80 buffer (TEKnova) using in-house protocols. The coding regions of all constructs and mutants were verified by DNA sequencing using MRC PPU DNA Sequencing and Services or Eurofins Genomics DNA sequencing services.

### Expression and purification of recombinant proteins

All E2, Ubiquitin and PCNA constructs were transformed into chemically competent BL21 *E. coli* strains and cultured in Miller LB broth (Merck) at 37°C until OD_600_ ~ 0.4 following which temperature was reduced to 16°C. At OD_600_ ~ 0.8 protein expression was induced by adding a final concentration of 200 μM (for Ube2T and Ube2D3) or 500 μM (for Ube2B, Ubiquitin and PCNA) Isopropyl-Beta-d-Thiogalactopyranoside (IPTG, Formedium) and allowed to proceed for a further 16–18h. Similar steps were followed for all RING domain constructs except the growth media was supplemented with 250 μM ZnCl^2^ and protein expression was induced at OD_600_ ~ 1.0 with a final concentration of 50 μM IPTG. All purification steps, except for ubiquitin, were performed at 4°C and completed within 24–36 hours of cell lysis.

Purification of all Ube2T variants and FANCL fragments are as described in (Hodson et al., 2011; Hodson et al., 2014). The respective affinity tags were cleaved using GST-3C protease (for Ube2T) and 6xHis-Ulp1 protease (for FANCL). Ube2B variants and PCNA were expressed and purified similar to Ube2T. The 6xHis tag was removed from Ube2B using 6xHis-TEV protease. The 6xHis-smt3-Rad18 and 6xHis-MBP-rat RNF4 RING-RING proteins was purified as described elsewhere (Huang et al., 2011; Plechanovova et al., 2011). The 6xHis-smt3 was retained for RAD18 and protein concentration for both proteins were estimated using SDS-PAGE and coomassie staining using known protein standards of similar size.

Baculovirus were generated using the Bac-to-Bac (Invitrogen) system and proteins were expressed for ~72h in baclovirus infected *Sf*21 cells. Cell pellets were suspended in lysis buffer (50 mM Tris-HCl pH8.0, 150 mM NaCl, 10 mM imidazole, 1 mM TCEP and 5% v/v glycerol with freshly added 2 mM MgCl^2^, EDTA-free Protease Inhibitor cocktail (Pierce) and Benzonase (Sigma Aldrich). *Sf*21 cells were lysed by homogeniser followed by sonication in an ice-bath. All lysates were clarified at 40,000 RCF for 45 min at 4°C and filtered. Proteins were bound to HisPur Ni-NTA Resin (Thermo Scientific) and washed extensively with lysis buffer with 500 mM NaCl. 6xHis-TEV-FANCI or His-3C-FANCD2 proteins were eluted by lysis buffer with 250 mM imidazole and lower NaCl for ion exchange chromatography (50 mM NaCl). Anion exchange for FANCI and FANCD2 were all performed using a 5 ml HiTRAP Q HP column (GE Life Sciences) using AKTA FPLC systems and eluted with a linear gradient (50 mM Tris pH8.0, 50–1000 mM NaCl, 5 mM Dithiothreitol (DTT) and 5% v/v glycerol). The proteins were further purified using SEC using a Superose6 10/300 GL column in 20 mM Tris pH8.0, 400 mM NaCl, 5 mM DTT and 5% v/v glycerol. Proteins were concentrated using 50,000 MWCO Amicon Ultra centrifugal filters (Merck) to before flash-frozen as single-use aliquots in the same buffer system.

### Preparation of Ub^IR800^ material

6xHis-smt3-SGCGSG-Ubiquitin material post affinity and gel-filtration chromatography was dialysed into 1x Phosphate buffered saline (PBS) pH7.4 with 0.5 mM EDTA. Protein concentration was estimates using SDS-PAGE and SimplyBlue (Invitrogen) staining using known protein standards of similar size. The dialysed material was reduced with a final concentration of 50 mM 2-Mercaptoethanol (Sigma Aldrich) for 1 hour at 37°C. The material was rapidly buffered exchanged into 1xPBS pH7.4 with 0.5 mM EDTA using 7K MWCO Zeba Spin Desalting Columns (Pierce) and immediately mixed, at 1:2 ratio, with DyLight 800 Maleimide (Life Technologies) dye solubilised in neat Dimethylformamide (DMF, Pierce). All subsequent steps were protected from direct light. The labelling reaction was allowed to proceed at 25°C for 8–10h, quenched using 50 mM 2-Mercaptoethanol and excess dye removed by extensive dialyses into 1x PBS buffer using 10,000 MWCO Spectra/Por membrane (Spectrum Labs). The labelled fusion protein was cleaved using 6xHis-Ulp1 protease,passed over Ni-NTA Resin (Thermo Scientific) to capture the protease and 6xHis-smt3 tag. The material was further purified using Superdex 75 10/300 GL in 50 mM Tris-HCl pH7.5, 150 mM NaCl. Fractions with labelled ubiquitin were pooled and protein concentration was estimated as above.

### Multi-turnover substrate ubiquitination assays

All reactions were carried out in 50 mM Tris-HCl, 100 mM NaCl, 1 mM TCEP, 2 mM MgCl^2^ and 4% v/v glycerol buffer system at pH7.6 and 30°C. Frozen protein aliquots were thawed on ice and reactions performed within 12 h of thaw. All reaction mixtures were performed on ice. The DNA co-factors for human substrates have been previously described (Longerich et al., 2014) and the following sequences were synthesised as 1 μmole duplexes and PAGE purified (IDT technologies).

Sense -

TTGATCAGAGGTCATTTGAATTCATGGCTTCGAGCTTCATGTAGAGTCGACGGTGCTGGGATCCAACATGTTTTCAATCTG

Antisense -

CAGATTGAAAACATGTTGGATCCCAGCACCGTCGACTCTACATGAAGCTCGAAGCCATGAATTCAAATGACCTCTGATCAA

End-point reactions (15 μL) contained 25 nM 6xHis-Ube1, 3 μM Ub^IR800^, 0.1 μM Ube2T and FANCL^UR^, 1 μM FANCI-FANCD2 complex or FANCI alone and 2 μM dsDNA. Reaction mixes were incubated on ice for 10 min to allow for substrate-DNA complex formation followed by addition of Adenosine triphosphate (ATP) at a final concentration of 5 mM. Reactions were terminated after 30 min with an equal amount of 2xLDS sample buffer (Pierce) containing 200 mM 2-Mercaptoethanol and boiled at 98°C for 3min. Subsequently 4 μL of boiled samples were resolved in 10 well Bolt 4–12% Bis-Tris Plus Gels (Invitrogen) using a 1x MOPS running buffer system (Invitrogen). Gels were resolved until the 25 kDa MW marker of All Blue Precision Plus protein standard (Bio-Rad) is at the bottom of the gel.

Gels were rinsed with water and scanned by direct fluorescence monitoring using Li-COR Odyssey CLX Infrared Imaging System. Time-course reactions (60 μL) for rate determination were setup as described above except the 0 min time-point was taken prior to addition of ATP, 8 μL sample at indicated time-points and terminated with equal amounts of 2xLDS reducing sample buffer. Samples were boiled together after last time-point and 4 μL were resolved in 17 well NuPAGE 4–12% Bis-Tris Gels (Invitrogen) as described above. Gels were scanned as before and analysed using ImageStudio software (Li-COR). A custom rectangle in 0 min lane was used for background subtraction. An identical shape area was used to quantify amount of product formed using trimmed signal intensities values. The data was exported into Microsoft Excel and plotted against time to determine rates in the linear reaction range. Finally rates were normalized to wildtype E2. Experiments were performed in triplicate and final rate graphs were plotted (Mean ± SD) in GraphPad Prism 7.

PCNA ubiquitination assays (20 μL) contained 25 nM 6xHis-Ube1, 20 μM Ub, 2 μM Ube2B, 9 μM 6xHisPCNA and the indicated amounts of 6xHis-smt3-Rad18. Post ATP addition (5 mM), reactions were allowed to proceed for 90 min and terminated with equal amounts of 2xLDS reducing sample buffer and boiled as before. Samples were diluted to 60 μL using 1xLDS reducing sample buffer. 3 μL (PCNA blot) and 10 μL (Ube2B and Rad18 blot) of the diluted sample was resolved in 15 well NuPAGE 4–12% Bis-Tris Gels (Invitrogen) and transferred using iBlot Western Blotting system and nitrocellulose transfer stacks (Invitrogen). Membranes were blocked using 1xPBS buffer containing 1% w/v BSA and 0.05% v/v Tween20. The respective membranes were incubated overnight at 4°C with anti-PCNA mouse monoclonal antibody (ab29, Abcam) at 0.4 ng/μL, anti-Ube2B rabbit polyclonal (10733–1-AP, Proteintech) at 0.15 ng/μL and anti-Rad18 rabbit polyclonal antibody (18333–1-AP) at 0.6 ng/μL. Blots were washed 3×15min with 1xPBS buffer with 0.05% v/v Tween20 and probed with IR800 labelled secondary antibodies (Li-COR) of the corresponding species at 0.1 ng/μL for 2 h at room-temperature. Blots were washed 3×15min with 1xPBS buffer with 0.05% v/v Tween20 and scanned by direct fluorescence monitoring using Li-COR Odyssey CLX Infrared Imaging System. Experiments were performed in duplicate to confirm consistency of results

### E2 charging, autoubiquitination and lysine discharge assays

All reactions were carried out in 50 mM Hepes, 100 mM NaCl, 1 mM TCEP, 2 mM MgCl^2^ and 4% v/v glycerol buffer system at pH7.6 and 30°C. Ube2T charging reactions (10 μL) contained 100 nM 6xHis-Ube1, 10 μM E2 and Ubiquitin. Reactions were commenced by addition of buffer or ATP at a final concentration of 5 mM. Reactions were terminated after 5 min with 5 μL of non-reducing 3xLDS sample buffer. 1 μg of Ube2T was resolved in 15 well NuPAGE 4–12% Bis-Tris Gels (Invitrogen). Gels were rinsed with water, stained with InstantBlue Coomassie stain. Gels were de-stained with water and scanned. Experiments were performed in duplicate to confirm consistency of results. Ube2T^1–152^ auto-ubiquitination reactions (60 μL) contained 100 nM 6xHis-Ube1, 10 μM E2 and 20 μM Ubiquitin. The 0 time-point was taken prior to ATP addition (5 mM final), 8 μL sample was taken at indicated time-points and terminated with 4 μL of 3xLDS reducing sample buffer. 0.8 μg of Ube2T^1–152^ was resolved in 15 well NuPAGE 12% Bis-Tris Gels (Invitrogen), stained and de-stained as before. Experiments were performed in triplicate to confirm consistency of results. For lysine discharge assays, charging reactions (50 μL) contained 120 nM 6xHis-Ube1, 12 μM E2 and 24 μM Ubiquitin and started with ATP (5 mM final). After 10 min at 30°C, 0.5 U of Apyrase (NEB) was added to arrest the reaction and left at room temperature for 3 min. The chase was initiated by adding 10 mM L-Lysine (Sigma Aldrich) to bring the final volume to 60 μL. Initial time-point sample (8 μL) was taken at 0.1 min and subsequent samples were taken at indicated time-points. Samples were terminated with 4 μL of 3xLDS non-reducing sample buffer. 0.9 μg of Ube2T^1–152^ was resolved in 15 well NuPAGE 12% Bis-Tris Gels (Invitrogen), stained and de-stained as before. Gels were scanned by direct fluorescence monitoring (700 nm λ) using Li-COR Odyssey CLX Infrared Imaging System. The intensities of product formed (E2 released) was obtained and converted protein amount (picomoles) using an E2 only serial dilution reference gel that was stained and de-stained in parallel. Absolute rates were determined by plotting product versus time in Microsoft Excel. Experiments were performed in duplicate and final rate graphs were plotted (Mean ± SD) in GraphPad Prism 7.

### Single-turnover substrate ubiquitination assays

All reactions were carried out in 50 mM Tris-HCl, 100 mM NaCl, 1 mM TCEP, 2 mM MgCl^2^ and 4% v/v glycerol buffer system at pH7.6. Charging reactions (30 μL) contained 10 nM 6xHis-Ube1, 10 μM E2 and 10 μM Ub^IR800^ and started with ATP (5 mM final). After 10 min at 30°C, 0.25 U of Apyrase was added to arrest the reaction and left at room temperature for 3 min. E2 charging efficiency was determined to be ~80%. Chase mixes (40 μL) containing 2 μM FANCI^K523R^-FANCD2-dsDNA complex, 0.2 μM E2~Ub^IR800^ and indicated amounts of FANCL^UR^ were incubated at 30°C. After 10 min, 5 μL of the reaction diluted 1- fold and terminated with 10 μL of 2xLDS non-reducing or reducing sample buffer. Samples with reducing agent were boiled at 98°C for 3min. 4 μL of each sample were resolved in 10 well Bolt 4–12% Bis-Tris Plus Gels (Invitrogen) until the 25 kDa MW marker is at the bottom of the gel. Gels were rinsed with water and scanned by direct fluorescence monitoring using Li-COR Odyssey CLX Infrared Imaging System.

### Isothermal titration calorimetry

ITC experiments were performed using MicroCal PEAQ-ITC (Malvern instruments). All experiments were performed at 20°C, in duplicate, using freshly prepared proteins within 2 days of the last purification step. Proteins were buffer exchanged using 7K MWCO Zeba Spin Desalting Columns (Pierce) into 100 mM Tris-HCl, 100 mM NaCl, 0.4 mM TCEP buffer at pH 8.0 that was filtered and thoroughly degassed. FANCL^UR^ (ranging 22 to 34 μM) and FANCL^R^ (~32 μM) was held in the cell, while Ube2T (ranging 400 to 600 μM) was present in the syringe. A total of 16 injections were carried out with the first injection of 0.3 μL over 0.6s followed by 15 injections of 1.5 μL over 3s. All injections were spaced by 120s with mix speed set at 500 rotations per minute. Each experiment was controlled by an identical E2 into buffer run to account for the heat of dilution. All data were fitted using a single-site binding model using MicroCal PEAQ-ITC analysis software.

### Residue Network Analysis

Residue Interaction Networks (RINs) were generated using the RIN generator webserver (http://protein.bio.unipd.it/ring/)(Piovesan et al., 2016). For this study, networks were generated for consecutive residues in individual chains using relaxed distance thresholds. Further, all atoms of a likely residue pair were considered when applying distance thresholds and subsequently all possible interactions outputted. Finally, water molecules and hetero atoms/ligands were omitted for the RIN. The output files were uploaded in Cytoscape (Shannon et al., 2003), and the RINs were compared using the RINalyzer app (Doncheva et al., 2011) using residue IDs as matching attribute. In case of Ube2T, two copies of the RING bound state were first individually compared to unbound state using a difference edge weight. The resulting RIN comparison matrices were merged into a collective network using the inbuilt merge network tool on Cytoscape.

## Author contributions

V.K.C. – Conceptualization, Methodology, Validation, Formal Analysis, Investigation, Resources, Writing - original draft preparation, review and editing, Visualization, Project Administration.

C.A. – Methodology, Validation, Resources, Writing - review.

R.T. – Resources.

H.W. – Conceptualization, Writing - review and editing, Supervision, Project Administration, Funding Acquisition

## Acknowledgements

This work was supported by the Medical Research Council (MRC grant number MC_UU_12016/12); the EMBO Young Investigator Programme to H.W.; the European Research Council (ERC-2015-CoG-681582 ICLUb consolidator grant to H.W. The pLou3 Rat RNF4 RING-RING DNA construct was a kind gift from Prof. Ron Hay, University of Dundee. The human PCNA cDNA template, was a kind gift from Dr Svend Petersen-Mahrt, European Institute of Oncology, Milan. The Human 6xHis-Ube1 protein material was a kind gift from Dr. Axel Knebel, MRC Protein Phosphorylation and Ubiquitylation Unit. All constructs are available on request from the MRC Protein Phosphorylation and Ubiquitylation Unit reagents Web page (http://mrcppureagents.dundee.ac.uk) or from the corresponding author.

## Conflict of interest

Authors declare no conflict of interest.

---

**Supplementary Table 1**. Edge profile of Ube2T allosteric network model depicted in Figure 4a.

**Supplementary Dataset S1**. Merged Ube2T intra-residue network.

**Supplementary Dataset S2**. Merged Ube2B intra-residue network.

**Supplementary Figure 1.**
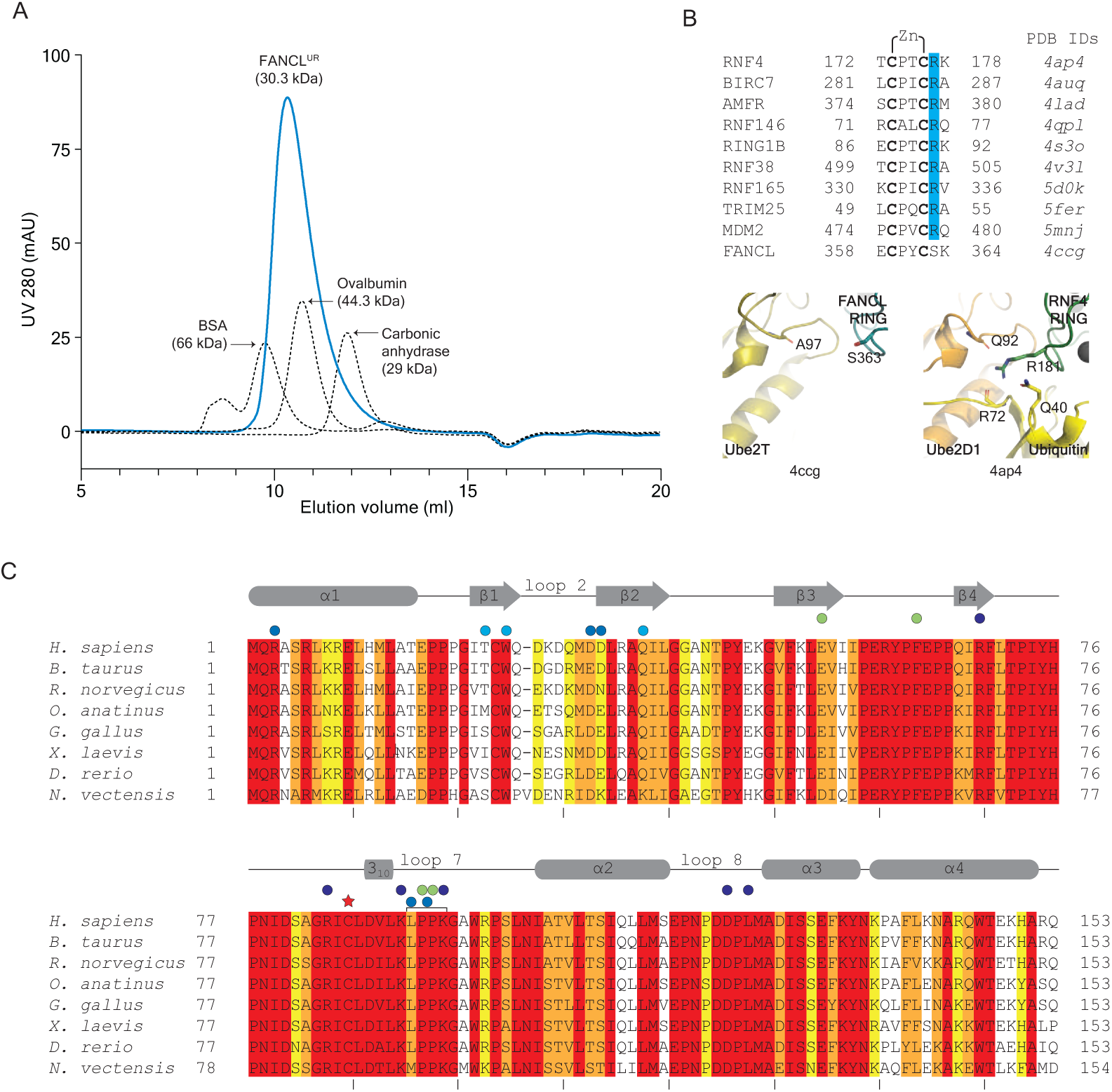
Solution and sequence profile of FANCL^UR^ with Ube2T sequence conservation. A Analytical size-exclusion chromatography of FANCL^UR^ using Superdex75 10/300 GL column (blue trace). Gel-filtration standards (dashed trace) with known elution profile were also run in same buffer system as FANCL^UR^ and the traces were overlaid. The FANCL^UR^ fragment resolves as a monomeric protein however, the moderate shift to an earlier elution volume suggests the central UBC-RWD and C-terminal RING domain adopt an extended conformation as observed in the fly FANCL structure (PDB ID 3k1l). B Sequencing alignment of the C-terminal regions of various RING domains that have been characterised to have a linchpin Arg residue (blue highlight). FANCL appears to lack such a residue at the analogous position. Structure of the RNF4 linchpin residue (Arg181, green cartoon, PDB ID 4ap4) shows how the linchpin contacts both the E2 (Ube2D1, orange cartoon) and donor ubiquitin (yellow cartoon). In contrast, the equivalent FANCL residue Ser363 would not stabilise such conformation. Zinc co-ordinating cysteine residues are in bold. PDB IDs of respective structures are listed alongside. Molecular figures prepared in PyMOL (Schrödinger, LLC). C Sequence conservation of the UBC fold between Ube2T homologs. Residues shaded in red to yellow to highlight conservation, where red corresponds to strict conservation. Depicted above the sequences (grey) are secondary structure elements of human Ube2T are based on PDB ID 1yh2. Red star indicates catalytic cysteine while coloured circles indicate human Ube2T residues mutated in this study; blue in Figure 2, cyan in Figure 3, purple in Figure 4 and green in Figure 5.

**Supplementary Figure 2.**
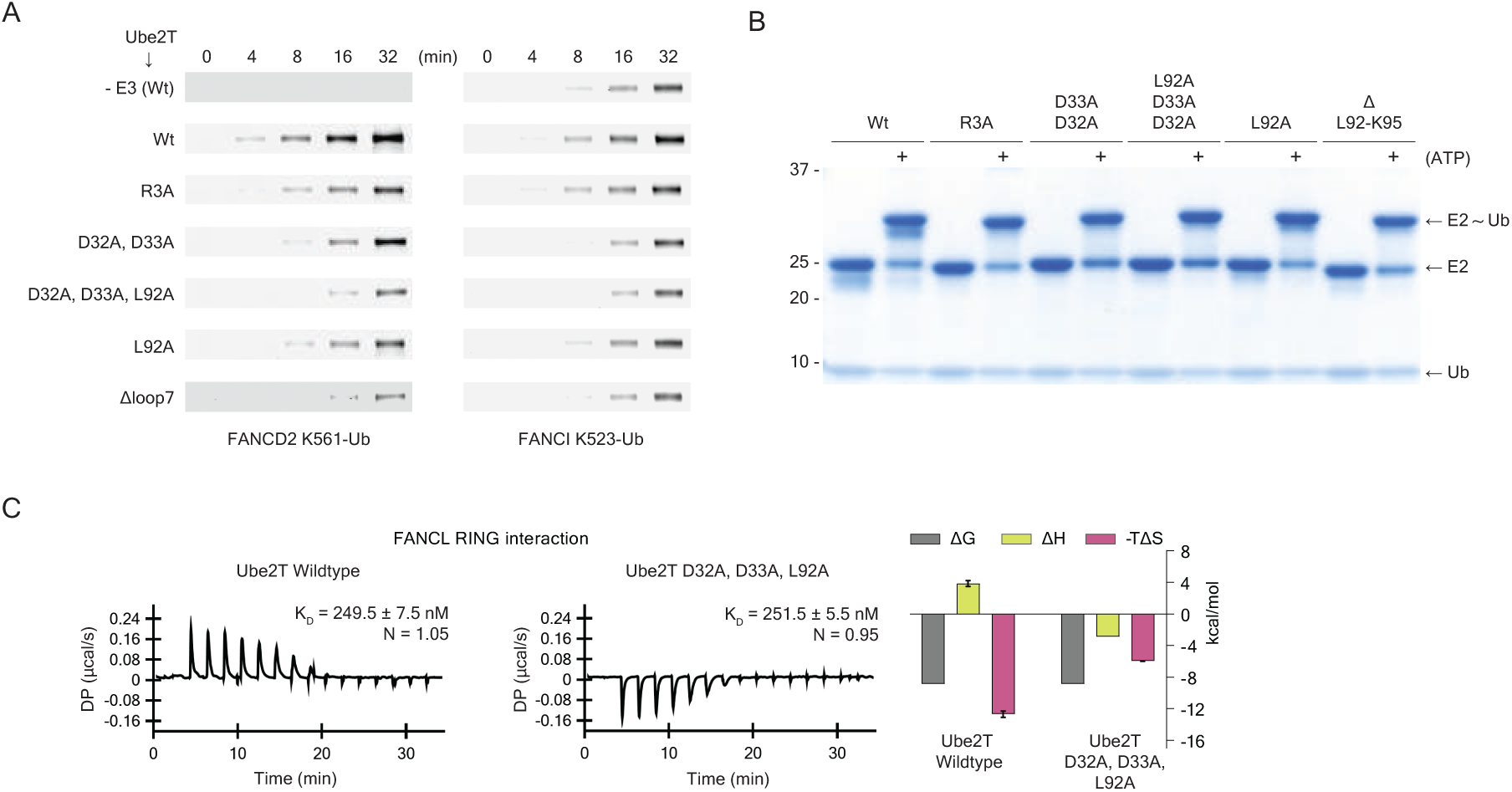
Activity and binding profile of Ube2T loop2 and loop7 residues. A Representative time-course ubiquitination assays with fluorescently labelled ubiquitin (Ub^IR800^) used to derive substrate ubiquitination rates depicted in Figure 2C. Substrate ubiquitination is analysed by direct fluorescence monitoring (Li-COR Odyssey CLX). B Ubiquitin charging assays of Ube2T loop2 and loop7 mutants show that the mutants do not affect E1-based E2~Ub thioester formation. C Thermodynamics of FANCL^R^ interaction with Ube2T wildtype (left) and Ube2T loop2-loop7 hybrid mutant (middle) shows similar contrast in binding enthalpy profile as observed with FANCL^UR^. Graphs (right) plotted as mean ± range (n=2).

**Supplementary Figure 3.**
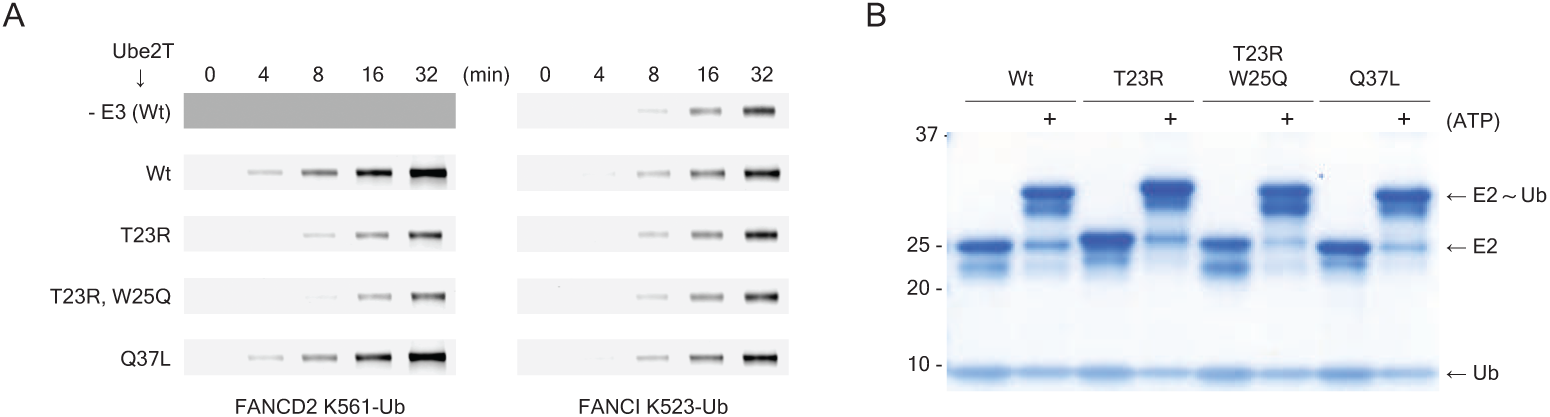
Activity profile of Ube2T backside residues. A Representative time-course ubiquitination assays with fluorescently labelled ubiquitin (Ub^IR800^) used to derive substrate ubiquitination rates depicted in Figure 3D. Substrate ubiquitination is analysed by direct fluorescence monitoring (Li-COR Odyssey CLX). B Ubiquitin charging assays of Ube2T backside mutants show that the mutants do not affect E1-based E2~Ub thioester formation.

**Supplementary Figure 4.**
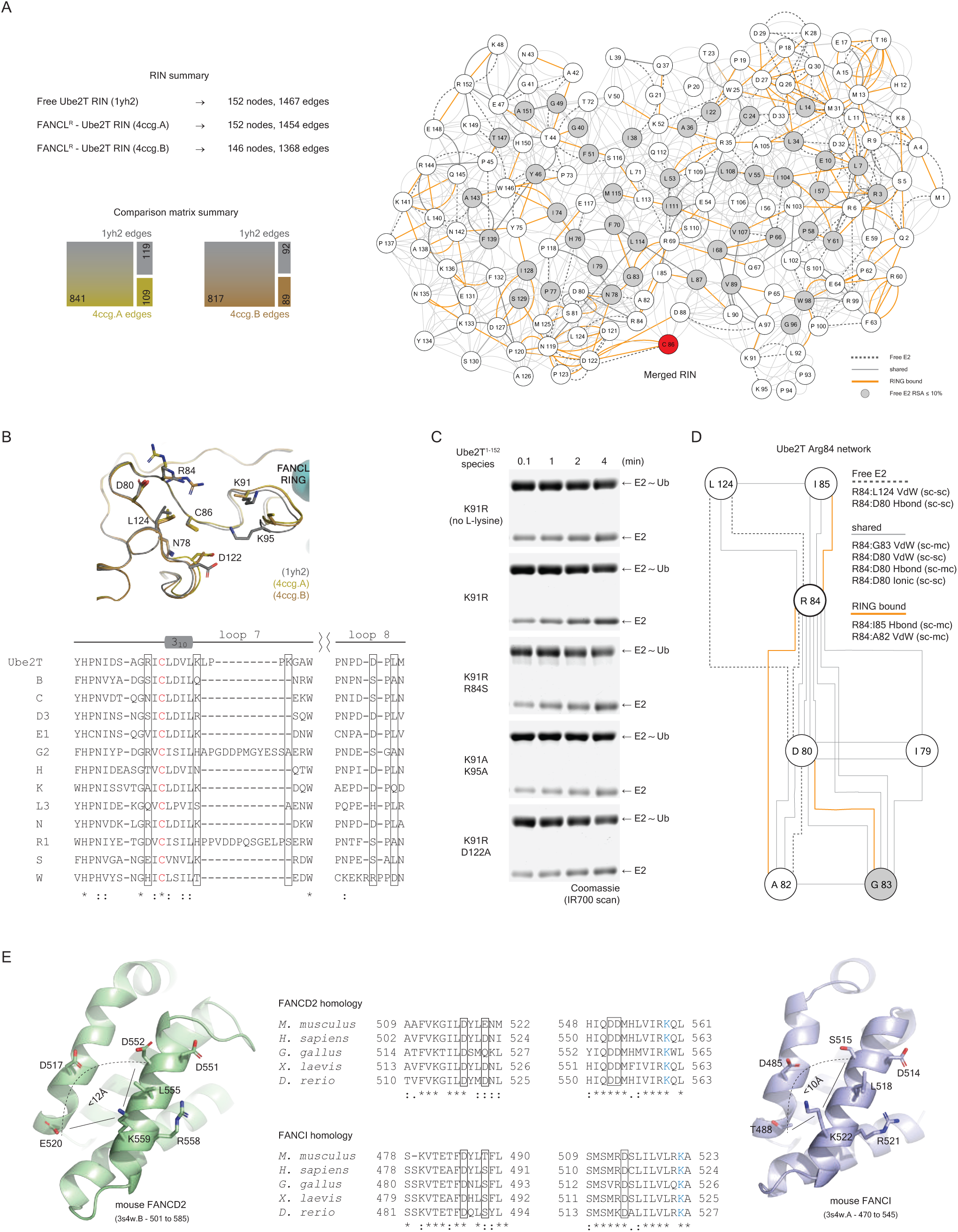
Ube2T residue network profile and characterization of network termini residues. A Summary of nodes and edges (top left) in the individual Ube2T residue interaction networks (RINs). Mosaic plots of edge distribution profile derived from RIN comparison (bottom left). Complete profile of edges and nodes obtained after merging the RIN comparison networks. Dashed and orange lines depict Ube2T edges in unbound and FANCL^R^ bound state respectively. Thin grey lines depict edges that are shared in both networks. Grey nodes have relative solvent accessibility of less than 10% in unbound Ube2T. Red node denotes the catalytic cysteine (Cys86). B Structural overlay (top) of unbound and RING bound Ube2T structures focussing on network termini residues surrounding the catalytic cysteine. Sequence alignment (bottom) of Ube2T catalytic site with various other human E2s reveal how the termini of Ube2T network (indicated by boxes) vary among the enzyme family. Catalytic cysteine is shown in red. Molecular figures prepared in PyMOL (Schrödinger, LLC). C Representative gels of lysine discharge assays plotted in Figure 4D. Coomassie stained gels are visualized by direct fluorescence monitoring in the red channel (Li-COR Odyssey CLX). D Total network profile of Arg84 residue present in Ube2T catalytic β-element. Listed alongside are edges that involve the Arg84 side chain. Edge depiction is same as in panel A. E Acidic/polar residues found proximal to target lysine on FANCD2 (left, green cartoon) and FANCI (right, blue cartoon) based on the mouse FANCI-FANCD2 complex structure (PDB ID 3s4w). Sequence alignment (middle) shows conservation of these acidic/polar residues (boxed). Respective target lysine is depicted in blue. Molecular figures prepared in PyMOL (Schrödinger, LLC).

**Supplementary Figure 5.**
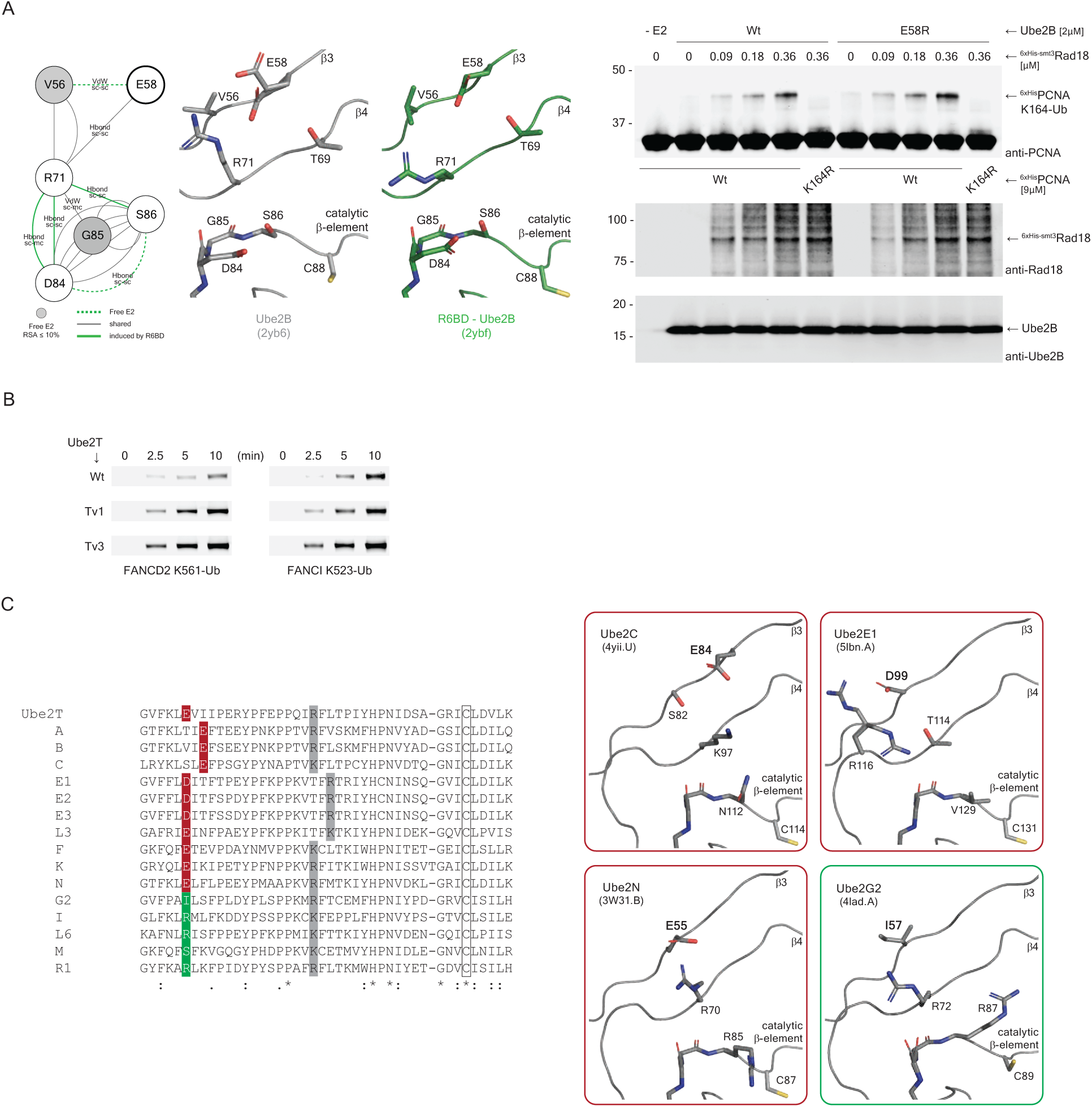
Activity profile of deregulated Ube2B and Ube2T variants. A Comparison of a network scheme (top left) with structures of unbound (top middle, PDB ID 2yb6) and Rad6 Binding Domain (R6BD) bound (top right, PDB ID 2ybf) Ube2B depicting the proposed gating and effector roles for Ube2B residues Glu58 (thick outline) and Arg71 respectively. Green dashed and green lines depict Ube2B edges in unbound and R6BD bound states respectively. Grey lines depict common connections in the two Ube2B states while grey nodes have relative solvent accessibility of less than 10% in unbound E2. Immunoblots of an end-point (90 min) multi-turnover PCNA ubiquitination assay (bottom) shows Ube2B with a permissive Glu54Arg gate is more sensitive to increasing amounts of Rad18 E3 in substrate ubiquitination. Control immunoblots anti-Rad18 (middle) and anti-Ube2B (bottom) show levels of E3 and E2 respectively. B Representative time-course ubiquitination assays with fluorescently labelled ubiquitin (Ub^IR800^) used to derive substrate ubiquitination rates depicted in Figure 5E. Substrate ubiquitination is analysed by direct fluorescence monitoring (Li-COR Odyssey CLX). C Structures of human E2s with the proposed β3 gating residue (bold label), restrictive in red boxes while permissive in green boxes. Depicted below is a sequence alignment of human E2s, restrictive gate highlighted in red while permissive gate in green. Highlighted in grey is the proposed effector residue linked with the gate. Ube2T and Ube2B sequences are aligned as reference. Highlighted in grey is the proposed effector residue liked with the gate. Catalytic cysteine is also indicated (clear box). Molecular figures prepared in PyMOL (Schrödinger, LLC).

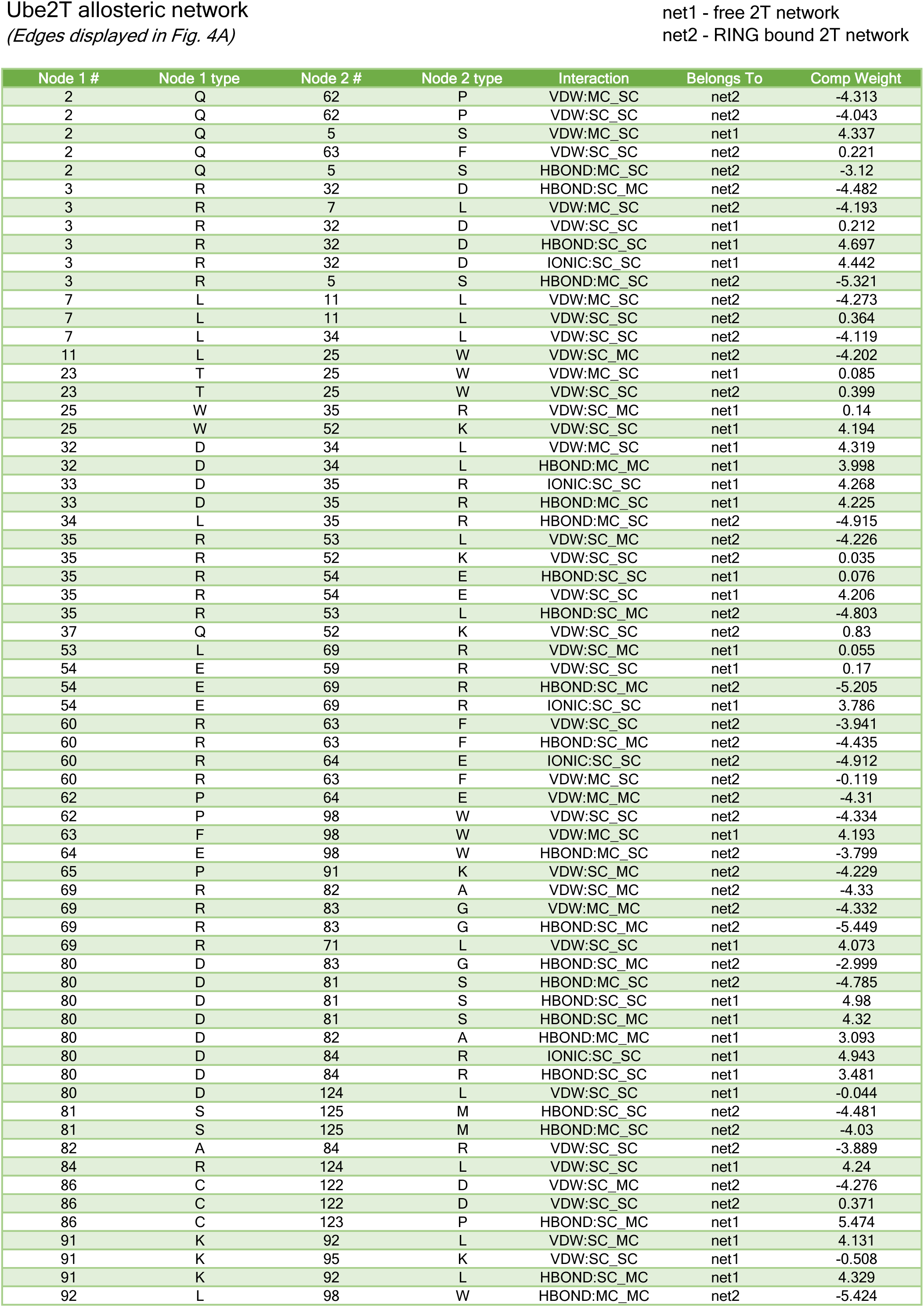
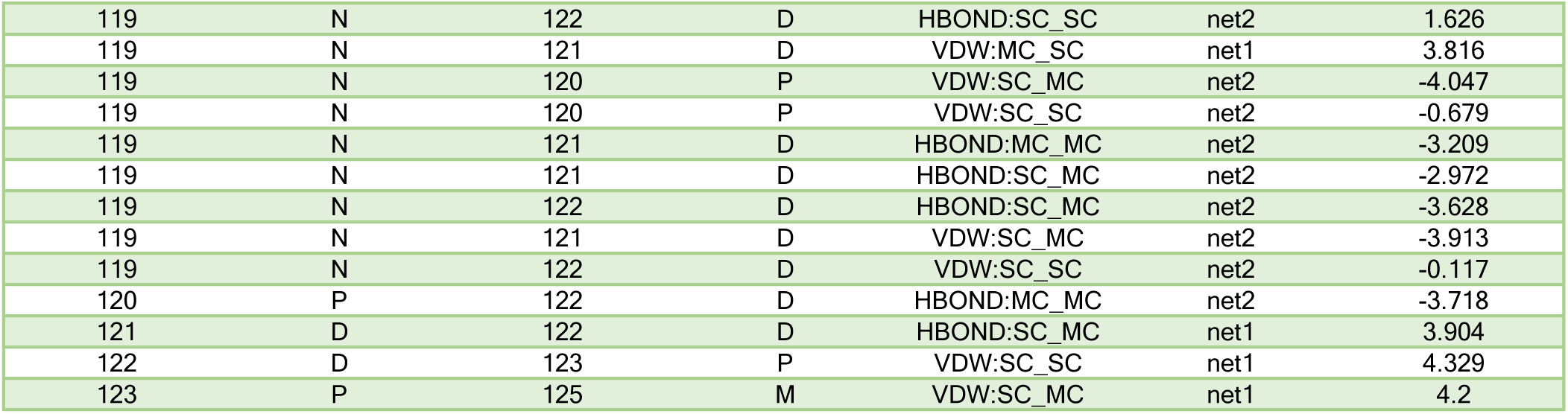

## References

Al-Hakim, A., Escribano-Diaz, C., Landry, M.C., O’Donnell, L., Panier, S., Szilard, R.K., and Durocher, D. (2010). The ubiquitous role of ubiquitin in the DNA damage response. DNA Repair (Amst) 9, 1229–1240.

Alpi, A.F., Pace, P.E., Babu, M.M., and Patel, K.J. (2008). Mechanistic insight into site-restricted monoubiquitination of FANCD2 by Ube2t, FANCL, and FANCI. Mol Cell 32, 767–777.

Alpi, A.F., and Patel, K.J. (2009). Monoubiquitylation in the Fanconi anemia DNA damage response pathway. DNA Repair (Amst) 8, 430–435.

Benirschke, R.C., Thompson, J.R., Nomine, Y., Wasielewski, E., Juranic, N., Macura, S., Hatakeyama, S., Nakayama, K.I., Botuyan, M.V., and Mer, G. (2010). Molecular basis for the association of human E4B U box ubiquitin ligase with E2-conjugating enzymes UbcH5c and Ubc4. Structure 18, 955–965.

Bentley, M.L., Corn, J.E., Dong, K.C., Phung, Q., Cheung, T.K., and Cochran, A.G. (2011). Recognition of UbcH5c and the nucleosome by the Bmi1/Ring1b ubiquitin ligase complex. EMBO J 30, 3285–3297.

Branigan, E., Plechanovova, A., Jaffray, E.G., Naismith, J.H., and Hay, R.T. (2015). Structural basis for the RING-catalyzed synthesis of K63-linked ubiquitin chains. Nat Struct Mol Biol 22, 597–602.

Brown, N.G., VanderLinden, R., Watson, E.R., Qiao, R., Grace, C.R., Yamaguchi, M., Weissmann, F., Frye, J.J., Dube, P., Ei Cho, S., et al. (2015). RING E3 mechanism for ubiquitin ligation to a disordered substrate visualized for human anaphase-promoting complex. Proc Natl Acad Sci U S A 112, 5272–5279.

Brown, N.G., Watson, E.R., Weissmann, F., Jarvis, M.A., VanderLinden, R., Grace, C.R.R., Frye, J.J., Qiao, R., Dube, P., Petzold, G., et al. (2014). Mechanism of polyubiquitination by human anaphase-promoting complex: RING repurposing for ubiquitin chain assembly. Mol Cell 56, 246–260.

Ceccaldi, R., Sarangi, P., and D’Andrea, A.D. (2016). The Fanconi anaemia pathway: new players and new functions. Nat Rev Mol Cell Biol 17, 337–349.

Chakrabarti, K.S., Li, J., Das, R., and Byrd, R.A. (2017). Conformational Dynamics and Allostery in E2:E3 Interactions Drive Ubiquitination: gp78 and Ube2g2. Structure 25, 794–805 e795.

Cole, A.R., Lewis, L.P., and Walden, H. (2010). The structure of the catalytic subunit FANCL of the Fanconi anemia core complex. Nat Struct Mol Biol 17, 294–298.

Das, R., Liang, Y.H., Mariano, J., Li, J., Huang, T., King, A., Tarasov, S.G., Weissman, A.M., Ji, X., and Byrd, R.A. (2013). Allosteric regulation of E2:E3 interactions promote a processive ubiquitination machine. EMBO J 32, 2504–2516.

Das, R., Mariano, J., Tsai, Y.C., Kalathur, R.C., Kostova, Z., Li, J., Tarasov, S.G., McFeeters, R.L., Altieri, A.S., Ji, X., et al. (2009). Allosteric activation of E2-RING finger-mediated ubiquitylation by a structurally defined specific E2-binding region of gp78. Mol Cell 34, 674–685.

Doncheva, N.T., Klein, K., Domingues, F.S., and Albrecht, M. (2011). Analyzing and visualizing residue networks of protein structures. Trends Biochem Sci 36, 179–182.

Dou, H., Buetow, L., Sibbet, G.J., Cameron, K., and Huang, D.T. (2012). BIRC7-E2 ubiquitin conjugate structure reveals the mechanism of ubiquitin transfer by a RING dimer. Nat Struct Mol Biol 19, 876–883.

Eddins, M.J., Carlile, C.M., Gomez, K.M., Pickart, C.M., and Wolberger, C. (2006). Mms2-Ubc13 covalently bound to ubiquitin reveals the structural basis of linkage-specific polyubiquitin chain formation. Nat Struct Mol Biol 13, 915–920.

Ernst, A., Avvakumov, G., Tong, J., Fan, Y., Zhao, Y., Alberts, P., Persaud, A., Walker, J.R., Neculai, A.M., Neculai, D., et al. (2013). A strategy for modulation of enzymes in the ubiquitin system. Science 339, 590–595.

Freemont, P.S. (2000). RING for destruction? Curr Biol 10, R84–87.

Gabrielsen, M., Buetow, L., Nakasone, M.A., Ahmed, S.F., Sibbet, G.J., Smith, B.O., Zhang, W., Sidhu, S.S., and Huang, D.T. (2017). A General Strategy for Discovery of Inhibitors and Activators of RING and U-box E3 Ligases with Ubiquitin Variants. Mol Cell 68, 456–470 e410.

Gallego, L.D., Ghodgaonkar Steger, M., Polyansky, A.A., Schubert, T., Zagrovic, B., Zheng, N., Clausen, T., Herzog, F., and Kohler, A. (2016). Structural mechanism for the recognition and ubiquitination of a single nucleosome residue by Rad6-Bre1. Proc Natl Acad Sci U S A 113, 10553–10558.

Garaycoechea, J.I., and Patel, K.J. (2014). Why does the bone marrow fail in Fanconi anemia? Blood 123, 26–34.

Garcia-Higuera, I., Taniguchi, T., Ganesan, S., Meyn, M.S., Timmers, C., Hejna, J., Grompe, M., and D’Andrea, A.D. (2001). Interaction of the Fanconi anemia proteins and BRCA1 in a common pathway. Mol Cell 7, 249–262.

Hewitt, W.M., Lountos, G.T., Zlotkowski, K., Dahlhauser, S.D., Saunders, L.B., Needle, D., Tropea, J.E., Zhan, C., Wei, G., Ma, B., et al. (2016). Insights Into the Allosteric Inhibition of the SUMO E2 Enzyme Ubc9. Angew Chem Int Ed Engl 55, 5703–5707.

Hibbert, R.G., Huang, A., Boelens, R., and Sixma, T.K. (2011). E3 ligase Rad18 promotes monoubiquitination rather than ubiquitin chain formation by E2 enzyme Rad6. Proc Natl Acad Sci U S A 108, 5590–5595.

Hira, A., Yoshida, K., Sato, K., Okuno, Y., Shiraishi, Y., Chiba, K., Tanaka, H., Miyano, S., Shimamoto, A., Tahara, H., et al. (2015). Mutations in the gene encoding the E2 conjugating enzyme UBE2T cause Fanconi anemia. Am J Hum Genet 96, 1001–1007.

Hochstrasser, M. (2009). Origin and function of ubiquitin-like proteins. Nature 458, 422–429.

Hodson, C., Cole, A.R., Lewis, L.P., Miles, J.A., Purkiss, A., and Walden, H. (2011). Structural analysis of human FANCL, the E3 ligase in the Fanconi anemia pathway. J Biol Chem 286, 32628–32637.

Hodson, C., Purkiss, A., Miles, J.A., and Walden, H. (2014). Structure of the human FANCL RING-Ube2T complex reveals determinants of cognate E3-E2 selection. Structure 22, 337–344.

Huang, A., Hibbert, R.G., de Jong, R.N., Das, D., Sixma, T.K., and Boelens, R. (2011). Symmetry and asymmetry of the RING-RING dimer of Rad18. J Mol Biol 410, 424–435.

Joo, W., Xu, G., Persky, N.S., Smogorzewska, A., Rudge, D.G., Buzovetsky, O., Elledge, S.J., and Pavletich, N.P. (2011). Structure of the FANCI-FANCD2 complex: insights into the Fanconi anemia DNA repair pathway. Science 333, 312–316.

Kelly, A., Wickliffe, K.E., Song, L., Fedrigo, I., and Rape, M. (2014). Ubiquitin chain elongation requires E3-dependent tracking of the emerging conjugate. Mol Cell 56, 232–245.

Kim, W., Bennett, E.J., Huttlin, E.L., Guo, A., Li, J., Possemato, A., Sowa, M.E., Rad, R., Rush, J., Comb, M.J., et al. (2011). Systematic and quantitative assessment of the ubiquitin-modified proteome. Mol Cell 44, 325–340.

Kottemann, M.C., and Smogorzewska, A. (2013). Fanconi anaemia and the repair of Watson and Crick DNA crosslinks. Nature 493, 356–363.

Kulathu, Y., and Komander, D. (2012). Atypical ubiquitylation - the unexplored world of polyubiquitin beyond Lys48 and Lys63 linkages. Nat Rev Mol Cell Biol 13, 508–523.

Li, S., Liang, Y.H., Mariano, J., Metzger, M.B., Stringer, D.K., Hristova, V.A., Li, J., Randazzo, P.A., Tsai, Y.C., Ji, X., et al. (2015). Insights into Ubiquitination from the Unique Clamp-like Binding of the RING E3 AO7 to the E2 UbcH5B. J Biol Chem 290, 30225–30239.

Li, W., Tu, D., Li, L., Wollert, T., Ghirlando, R., Brunger, A.T., and Ye, Y. (2009). Mechanistic insights into active site-associated polyubiquitination by the ubiquitin-conjugating enzyme Ube2g2. Proc Natl Acad Sci U S A 106, 3722–3727.

Liang, C.C., Li, Z., Lopez-Martinez, D., Nicholson, W.V., Venien-Bryan, C., and Cohn, M.A. (2016). The FANCD2-FANCI complex is recruited to DNA interstrand crosslinks before monoubiquitination of FANCD2. Nat Commun 7, 12124.

Liu, W., Shang, Y., Zeng, Y., Liu, C., Li, Y., Zhai, L., Wang, P., Lou, J., Xu, P., Ye, Y., et al. (2014). Dimeric Ube2g2 simultaneously engages donor and acceptor ubiquitins to form Lys48-linked ubiquitin chains. EMBO J 33, 46–61.

Longerich, S., Kwon, Y., Tsai, M.S., Hlaing, A.S., Kupfer, G.M., and Sung, P. (2014). Regulation of FANCD2 and FANCI monoubiquitination by their interaction and by DNA. Nucleic Acids Res 42, 5657–5670.

Machida, Y.J., Machida, Y., Chen, Y., Gurtan, A.M., Kupfer, G.M., D’Andrea, A.D., and Dutta, A. (2006). UBE2T is the E2 in the Fanconi anemia pathway and undergoes negative autoregulation. Mol Cell 23, 589–596.

Mattiroli, F., Uckelmann, M., Sahtoe, D.D., van Dijk, W.J., and Sixma, T.K. (2014). The nucleosome acidic patch plays a critical role in RNF168-dependent ubiquitination of histone H2A. Nat Commun 5, 3291.

Mattiroli, F., Vissers, J.H., van Dijk, W.J., Ikpa, P., Citterio, E., Vermeulen, W., Marteijn, J.A., and Sixma, T.K. (2012). RNF168 ubiquitinates K13-15 on H2A/H2AX to drive DNA damage signaling. Cell 150, 1182–1195.

McGinty, R.K., Henrici, R.C., and Tan, S. (2014). Crystal structure of the PRC1 ubiquitylation module bound to the nucleosome. Nature 514, 591–596.

Meetei, A.R., de Winter, J.P., Medhurst, A.L., Wallisch, M., Waisfisz, Q., van de Vrugt, H.J., Oostra, A.B., Yan, Z., Ling, C., Bishop, C.E., et al. (2003). A novel ubiquitin ligase is deficient in Fanconi anemia. Nat Genet 35, 165–170.

Metzger, M.B., Liang, Y.H., Das, R., Mariano, J., Li, S., Li, J., Kostova, Z., Byrd, R.A., Ji, X., and Weissman, A.M. (2013). A structurally unique E2-binding domain activates ubiquitination by the ERAD E2, Ubc7p, through multiple mechanisms. Mol Cell 50, 516–527.

Metzger, M.B., Pruneda, J.N., Klevit, R.E., and Weissman, A.M. (2014). RING-type E3 ligases: master manipulators of E2 ubiquitin-conjugating enzymes and ubiquitination. Biochim Biophys Acta 1843, 47–60.

Middleton, A.J., and Day, C.L. (2015). The molecular basis of lysine 48 ubiquitin chain synthesis by Ube2K. Sci Rep 5, 16793.

Morreale, F.E., Bortoluzzi, A., Chaugule, V.K., Arkinson, C., Walden, H., and Ciulli, A. (2017). Allosteric Targeting of the Fanconi Anemia Ubiquitin-Conjugating Enzyme Ube2T by Fragment Screening. J Med Chem 60, 4093–4098.

Nijman, S.M., Huang, T.T., Dirac, A.M., Brummelkamp, T.R., Kerkhoven, R.M., D’Andrea, A.D., and Bernards, R. (2005). The deubiquitinating enzyme USP1 regulates the Fanconi anemia pathway. Mol Cell 17, 331–339.

Ozkan, E., Yu, H., and Deisenhofer, J. (2005). Mechanistic insight into the allosteric activation of a ubiquitin-conjugating enzyme by RING-type ubiquitin ligases. Proc Natl Acad Sci U S A 102, 18890–18895.

Petroski, M.D., and Deshaies, R.J. (2005). Mechanism of lysine 48-linked ubiquitin-chain synthesis by the cullin-RING ubiquitin-ligase complex SCF-Cdc34. Cell 123, 1107–1120.

Pickart, C.M. (2001). Mechanisms underlying ubiquitination. Annu Rev Biochem 70, 503–533.

Piovesan, D., Minervini, G., and Tosatto, S.C. (2016). The RING 2.0 web server for high quality residue interaction networks. Nucleic Acids Res 44, W367–374.

Plechanovova, A., Jaffray, E.G., McMahon, S.A., Johnson, K.A., Navratilova, I., Naismith, J.H., and Hay, R.T. (2011). Mechanism of ubiquitylation by dimeric RING ligase RNF4. Nat Struct Mol Biol 18, 1052–1059.

Plechanovova, A., Jaffray, E.G., Tatham, M.H., Naismith, J.H., and Hay, R.T. (2012). Structure of a RING E3 ligase and ubiquitin-loaded E2 primed for catalysis. Nature 489, 115–120.

Pruneda, J.N., Littlefield, P.J., Soss, S.E., Nordquist, K.A., Chazin, W.J., Brzovic, P.S., and Klevit, R.E. (2012). Structure of an E3:E2~Ub complex reveals an allosteric mechanism shared among RING/U-box ligases. Mol Cell 47, 933–942.

Rajendra, E., Oestergaard, V.H., Langevin, F., Wang, M., Dornan, G.L., Patel, K.J., and Passmore, L.A. (2014). The genetic and biochemical basis of FANCD2 monoubiquitination. Mol Cell 54, 858–869.

Rickman, K.A., Lach, F.P., Abhyankar, A., Donovan, F.X., Sanborn, E.M., Kennedy, J.A., Sougnez, C., Gabriel, S.B., Elemento, O., Chandrasekharappa, S.C., et al. (2015). Deficiency of UBE2T, the E2 Ubiquitin Ligase Necessary for FANCD2 and FANCI Ubiquitination, Causes FA-T Subtype of Fanconi Anemia. Cell Rep 12, 35–41.

Rodrigo-Brenni, M.C., Foster, S.A., and Morgan, D.O. (2010). Catalysis of lysine 48-specific ubiquitin chain assembly by residues in E2 and ubiquitin. Mol Cell 39, 548–559.

Rodrigo-Brenni, M.C., and Morgan, D.O. (2007). Sequential E2s drive polyubiquitin chain assembly on APC targets. Cell 130, 127–139.

Saha, A., Lewis, S., Kleiger, G., Kuhlman, B., and Deshaies, R.J. (2011). Essential role for ubiquitin-ubiquitin-conjugating enzyme interaction in ubiquitin discharge from Cdc34 to substrate. Mol Cell 42, 75–83.

Sareen, A., Chaudhury, I., Adams, N., and Sobeck, A. (2012). Fanconi anemia proteins FANCD2 and FANCI exhibit different DNA damage responses during S-phase. Nucleic Acids Res 40, 8425–8439.

Sato, K., Toda, K., Ishiai, M., Takata, M., and Kurumizaka, H. (2012). DNA robustly stimulates FANCD2 monoubiquitylation in the complex with FANCI. Nucleic Acids Res 40, 4553–4561.

Shannon, P., Markiel, A., Ozier, O., Baliga, N.S., Wang, J.T., Ramage, D., Amin, N., Schwikowski, B., and Ideker, T. (2003). Cytoscape: a software environment for integrated models of biomolecular interaction networks. Genome Res 13, 2498–2504.

Sheng, Y., Hong, J.H., Doherty, R., Srikumar, T., Shloush, J., Avvakumov, G.V., Walker, J.R., Xue, S., Neculai, D., Wan, J.W., et al. (2012). A human ubiquitin conjugating enzyme (E2)-HECT E3 ligase structure-function screen. Mol Cell Proteomics 11, 329–341.

Sims, A.E., Spiteri, E., Sims, R.J., 3rd, Arita, A.G., Lach, F.P., Landers, T., Wurm, M., Freund, M., Neveling, K., Hanenberg, H., et al. (2007). FANCI is a second monoubiquitinated member of the Fanconi anemia pathway. Nat Struct Mol Biol 14, 564–567.

Smogorzewska, A., Matsuoka, S., Vinciguerra, P., McDonald, E.R., 3rd, Hurov, K.E., Luo, J., Ballif, B.A., Gygi, S.P., Hofmann, K., D’Andrea, A.D., et al. (2007). Identification of the FANCI protein, a monoubiquitinated FANCD2 paralog required for DNA repair. Cell 129, 289–301.

Stewart, M.D., Ritterhoff, T., Klevit, R.E., and Brzovic, P.S. (2016). E2 enzymes: more than just middle men. Cell Res 26, 423–440.

Sugahara, R., Mon, H., Lee, J.M., and Kusakabe, T. (2012). Monoubiquitination-dependent chromatin loading of FancD2 in silkworms, a species lacking the FA core complex. Gene 501, 180–187.

Swuec, P., Renault, L., Borg, A., Shah, F., Murphy, V.J., van Twest, S., Snijders, A.P., Deans, A.J., and Costa, A. (2017). The FA Core Complex Contains a Homo-dimeric Catalytic Module for the Symmetric Mono-ubiquitination of FANCI-FANCD2. Cell Rep 18, 611–623.

Turco, E., Gallego, L.D., Schneider, M., and Kohler, A. (2015). Monoubiquitination of histone H2B is intrinsic to the Bre1 RING domain-Rad6 interaction and augmented by a second Rad6-binding site on Bre1. J Biol Chem 290, 5298–5310.

Uckelmann, M., and Sixma, T.K. (2017). Histone ubiquitination in the DNA damage response. DNA Repair (Amst) 56, 92–101.

Udeshi, N.D., Svinkina, T., Mertins, P., Kuhn, E., Mani, D.R., Qiao, J.W., and Carr, S.A. (2013). Refined preparation and use of anti-diglycine remnant (K-epsilon-GG) antibody enables routine quantification of 10,000s of ubiquitination sites in single proteomics experiments. Mol Cell Proteomics 12, 825–831.

Ulrich, H.D., and Walden, H. (2010). Ubiquitin signalling in DNA replication and repair. Nat Rev Mol Cell Biol 11, 479–489.

Valimberti, I., Tiberti, M., Lambrughi, M., Sarcevic, B., and Papaleo, E. (2015). E2 superfamily of ubiquitin-conjugating enzymes: constitutively active or activated through phosphorylation in the catalytic cleft. Sci Rep 5, 14849.

van den Ent, F., and Lowe, J. (2006). RF cloning: a restriction-free method for inserting target genes into plasmids. J Biochem Biophys Methods 67, 67–74.

van Twest, S., Murphy, V.J., Hodson, C., Tan, W., Swuec, P., O’Rourke, J.J., Heierhorst, J., Crismani, W., and Deans, A.J. (2017). Mechanism of Ubiquitination and Deubiquitination in the Fanconi Anemia Pathway. Mol Cell 65, 247–259.

Virts, E.L., Jankowska, A., Mackay, C., Glaas, M.F., Wiek, C., Kelich, S.L., Lottmann, N., Kennedy, F.M., Marchal, C., Lehnert, E., et al. (2015). AluY-mediated germline deletion, duplication and somatic stem cell reversion in UBE2T defines a new subtype of Fanconi anemia. Hum Mol Genet 24, 5093–5108.

Walden, H., and Deans, A.J. (2014). The Fanconi anemia DNA repair pathway: structural and functional insights into a complex disorder. Annu Rev Biophys 43, 257–278.

Wickliffe, K.E., Lorenz, S., Wemmer, D.E., Kuriyan, J., and Rape, M. (2011). The mechanism of linkage-specific ubiquitin chain elongation by a single-subunit E2. Cell 144, 769–781.

Windheim, M., Peggie, M., and Cohen, P. (2008). Two different classes of E2 ubiquitin-conjugating enzymes are required for the mono-ubiquitination of proteins and elongation by polyubiquitin chains with a specific topology. Biochem J 409, 723–729.

Yunus, A.A., and Lima, C.D. (2006). Lysine activation and functional analysis of E2-mediated conjugation in the SUMO pathway. Nat Struct Mol Biol 13, 491–499.

Zhang, W., Wu, K.P., Sartori, M.A., Kamadurai, H.B., Ordureau, A., Jiang, C., Mercredi, P.Y., Murchie, R., Hu, J., Persaud, A., et al. (2016). System-Wide Modulation of HECT E3 Ligases with Selective Ubiquitin Variant Probes. Mol Cell 62, 121–136.

Zhang, X.Y., Langenick, J., Traynor, D., Babu, M.M., Kay, R.R., and Patel, K.J. (2009). Xpf and not the Fanconi anaemia proteins or Rev3 accounts for the extreme resistance to cisplatin in Dictyostelium discoideum. PLoS Genet 5, e1000645.

